# Brg1 controls neurosensory cell fate commitment and differentiation in the mammalian inner ear

**DOI:** 10.1101/434159

**Authors:** Jinshu Xu, Jun Li, Yong-Hwee Eddie Loh, Ting Zhang, Huihui Jiang, Bernd Fritzsch, Aarthi Ramakrishnan, Li Shen, Pin-Xian Xu

## Abstract

Otic ectoderm gives rise to almost all cell types of the inner ear; however, the mechanisms that link transcription factors, chromatin, lineage commitment and differentiation capacity are largely unknown. Here we show that Brg1 chromatin-remodeling factor is required for specifying neurosensory lineage in the otocyst and for inducing hair and supporting cell fates in the cochlear sensory epithelium. Brg1 interacts with the critical neurosensory-specific transcription factors Eya1/Six1, both of which simultaneously interact with BAF60a or BAF60c. Chromatin immunoprecipitation-sequencing (ChIP-seq) and ChIP assays demonstrate Brg1 association with discrete regulatory elements at the *Eya1* and *Six1* loci. *Brg1*-deficiency leads to markedly decreased Brg1 binding at these elements and loss of *Eya1* and *Six1* expression. Furthermore, ChIP-seq reveals Brg1-bound promoter-proximal and distal regions near genes essential for inner ear morphogenesis and cochlear sensory epithelium development. These findings uncover essential functions for chromatin-remodeling in the activation of neurosensory fates during inner ear development.

## Introduction

The mammalian inner ear uses sensory hair cells (HCs) for mechanotransduction of vestibular and auditory stimuli and transmits this information to the brain via sensory neurons. Precursor cells for sensory cells and neurons are specified in the epithelium of the otocyst, which develops from the otic placode – thickened ectoderm that forms adjacent to the hindbrain at ~E8.5 and differentiates to form all inner ear structures. Neurosensory domain within the ventral region of the otic ectoderm is defined by transcription factors (TFs) Eya1 and Six1. A subpopulation of their daughter cells is induced to become neuroblasts from ~E9.0-9.25, which is marked by the basic helix-loop-helix (bHLH) TF Neurog1. These progenitors proliferate and differentiate to become Neurod1^+^ and then delaminate into the mesenchyme to aggregate and form the spiral and vestibular ganglion. In contrast, the prosensory progenitors continue to divide and expand within a common prosensory primordium until the commitment of sensory HC fate, which is marked by the bHLH protein Atoh1 from E12.5 in the vestibule and E14.5 in the cochlea. Sox2 is a crucial TF necessary for prosensory progenitor specification (Kiernan et al., 2005). We have shown that the transcription coactivator Eya1 and the homeodomain protein Six1 form a key transcriptional complex as neuronal or sensory HC determinants by demonstrating that expression of these two proteins can convert cochlear nonsensory epithelial GER (greater epithelial ridge) cells to a hair or neuronal cell fate through interaction with different partner proteins. Eya1-Six1 interacts with Sox2 to cooperatively activate an Atoh1-regulatory network for HC fate induction (Ahmed et al., 2012a) or with Brg1-BAFs (Brg1/Brm-associated factors) chromatin-remodeling proteins of the SWI/SNF (Switch/Sucrose non-fermentable) family to activate a Neurog1-Neurod1 regulatory network for neuronal differentiation (Ahmed et al., 2012b). Mutations in these factors cause congenital sensorineural hearing loss in humans or deafness in mice (Abdelhak et al., 1997; Hagstrom et al., 2005; Ozaki et al., 2004; Ruf et al., 2004; Xu et al., 1999; Zheng et al., 2003). However, the mechanisms that control the expression of *Eya1*, *Six1* and *Sox2* during acquisition of distinct cell fates are not understood. Furthermore, the SWI/SNF family plays a vital role in facilitating binding of specific TFs to nucleosomal DNA, yet no studies have been performed to directly address its roles in neurosensory cell fate specification and differentiation during inner ear development.

The SWI/SNF complex consists of multiple general and facultative subunits, whose activity requires Brg1 or the related Brm as ATPase (Ho and Crabtree, 2010). While *Brm*-null mice are viable and normal (Muchardt et al., 1998), deletion of *Brg1* causes early peri-implantation lethality (Bultman et al., 2000). Brg1 has been linked to progenitor cell proliferation, differentiation and survival in a variety of organs, including the heart, central nervous system, muscle, blood cells, and immune system (see reviews (Hargreaves and Crabtree, 2011; Ho and Crabtree, 2010). Since Eya1-Six1 interacts with the Brg1-BAFs, we conditionally deleted the central catalytic subunit Brg1 in *Eya1*-expressing cells at different developmental stages to investigate the function of the SWI/SNF complex during inner ear morphogenesis. Here we show that Brg1 is necessary for specifying both neuronal and sensory fates and is required for the expression of *Eya1* and *Six1*, both of which act upstream of *Sox2*. In the developing cochlea, Brg1 is not only necessary for prosensory domain specification but also for precursor cell differentiation into either HCs or supporting cells (SCs). Co-immunoprecipitation (coIP) shows a selective physical association of Eya1 and Six1 with the variants a and c but not b of the BAF60 structural subunit in the cochlea. Chromatin immunoprecipitation followed by deep sequencing (ChIP-seq) on E13.5 cochleae reveals Brg1-enriched regions both proximal to and distal from the promoters for genes essential for cochlea morphogenesis including *Eya1*, *Six1*, *Sox2*, *p27*^*Kip1*^ and *Gata3* loci. ChIP assays confirm co-occupancy of BAF60a/c at the Brg1-enriched regulatory elements at these loci. Conditional deletion of Brg1 results in significantly reduced Brg1 binding to these elements and loss of expression of these genes in the inner ear. Together, these results demonstrate that Brg1-based BAF complexes coordinate multiple processes of neurosensory lineage activation and differentiation and provide insight into how Brg1-BAFs act to promote lineage-specific properties in the inner ear.

## Results

### Brg1 controls specification of neurosensory progenitors in the otocyst

Brg1 is widely expressed in the otic placode and otocyst and in the delaminating neurons at E8.5-10.5 (Figure 1A,B). In the developing cochlea, Brg1 is highly expressed in the prosensory primordium marked by Sox2 (Figure 1C) and in differentiating HCs and SCs within the organ of Corti at P0 (Figure 1D). Strong Brg1 expression was also observed in LER (lesser epithelial ridge) and SGNs (spiral ganglion neurons) (Figures 1C,D and Figure 1-figure supplement 1A). To study its role in neurosensory cell development during inner ear morphogenesis, *Brg1*^*fl/fl*^ mice (Sumi-Ichinose et al., 1997) were bred with *Eya1*^*CreERT2*^ to generate *Eya1*^*CreERT2*^*;Brg1*^*fl/fl*^ (*Brg1*^*Cko/Cko*^ or CKO) mice with specific deletion of Brg1 in Eya1^+^ cells. Anti-Brg1 staining confirmed selective ablation in the ventral otocyst where Eya1 is normally expressed (Figure 1-figure supplement 1B-E). *Brg1*^*Cko/Cko*^ mice displayed a considerably smaller otocyst (*n*=6) than in their wild-type littermates and an absence of the VII-VIIIth ganglionic complex (Figure 1E). Examination of inner ears at E15.5 showed a degenerated cavity-like structure without neurosensory cells in *Brg1*^*Cko/Cko*^ (*n*=6, Figure 1F). These data indicate that Brg1 has an essential role in inner ear neurosensory structure formation.

**Figure 1.**
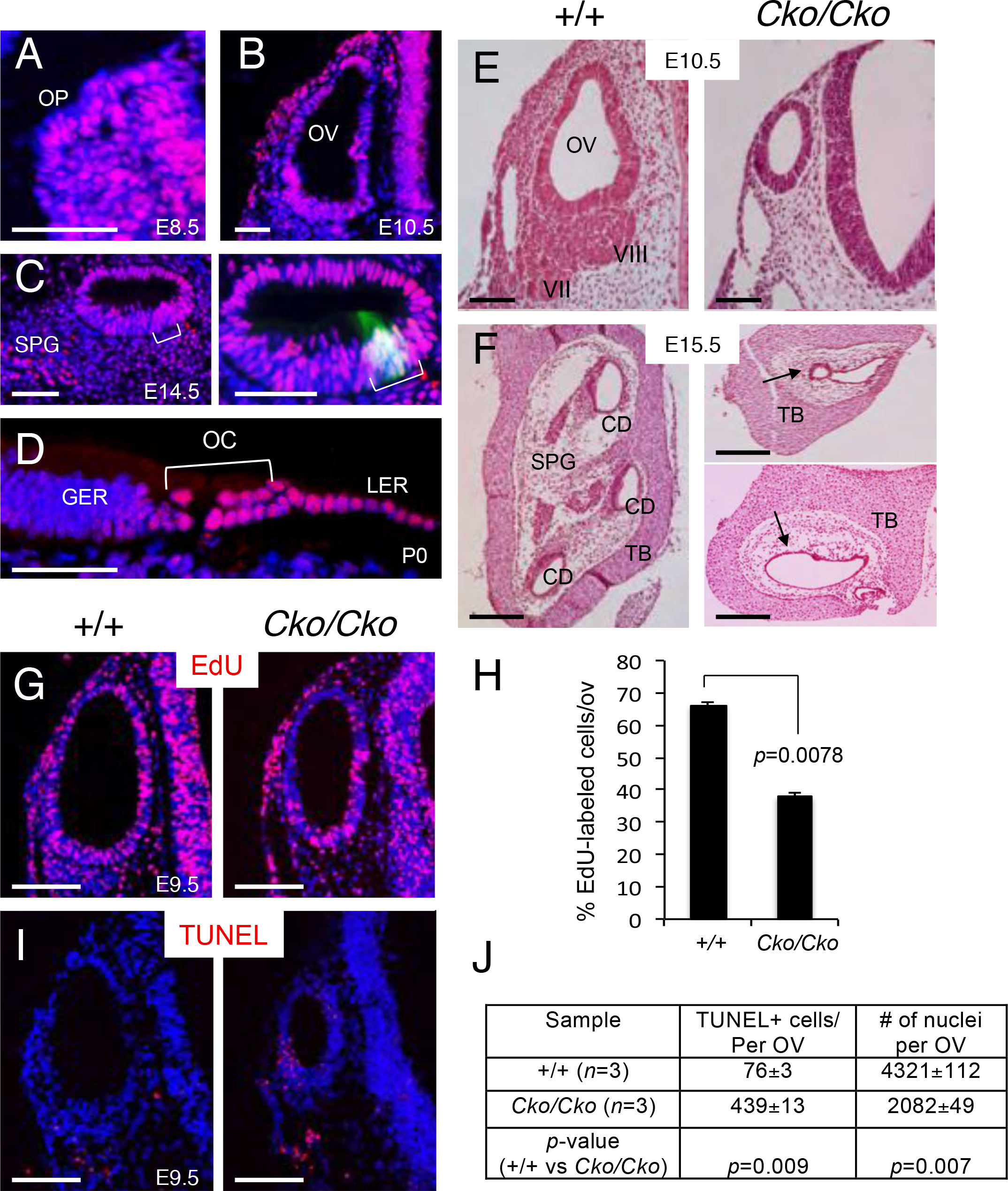
Deletion of *Brg1* using *Eya1*^*CreERT2*^ in the otic placode results in growth arrest and lack of inner ear structures. Tam was given from E8.5-9.0. (A-D) Anti-Brg1 staining in otic placode (A), otocyst (B), and developing cochlea (C,D). (E) H&E stained otocyst of wild-type and *Eya1*^*CreER*^*;Brg1*^*fl/fl*^ (*Cko/Cko*) littermates. (F) H&E stained inner ear sections of E15.5 wild-type and *Cko/Cko* littermates. Arrows indicate cavity-like structures. (G) EdU and (I) TUNEL assays. (H,J) Statistic analysis of EdU^+^ or TUENL^+^ cells from each otocyst. Data refer to the average of 3 embryos per genotype. *P*-value was calculated for +/+ and *Brg1*^*Cko/Cko*^ using Two-tailed Student’s *t*-test. Abb.: CD, cochlear duct; OC, organ of Corti; OV, otic vesicle; SPG, spiral ganglion; TB, temple bone. Scale bars: 50 μm (A-E, G,I), 200 μm (F).

Analysis of EdU (5-ethynyl-2’-deoxyuridine) incorporation (chased for 2 hours before harvest) revealed a reduction in proliferation rate in the mutant to ~57% of that in wild-type littermates (37.9±0.4% vs 66.1±0.3% per otocyst; *n*=3, Figure 1G,H). TUNEL assay showed increased abnormal cell apoptosis in the mutant (*n*=3, Figure 1I,J). Thus, the growth arrest and degeneration of the otocyst associated with Brg1-deficiency is due to defective cell proliferation and abnormal apoptosis.

To understand the molecular basis for the otic defects, we performed in situ hybridization (ISH) to characterize genes crucial for neurosensory cell fate specification. *Eya1*, *Six1* and *Sox2* are expressed in the ventral region of the otocyst from which all sensory organs develop (Figure 2A-E), whereas the expression of all three genes was not observed in the CKO littermates (*n*=6). *Dlx5*, which is expressed in the dorsolateral region of the otocyst (*n*=6, Figure 2F), was expressed in the ventral region of the CKO otocyst. This indicates that in the absence of Eya1-Six1-Sox2-promoted cell fate, the otic epithelium is filled with *Dlx5*-positive cells, suggesting acquisition of a non-sensory fate or expansion of the non-sensory area.

**Figure 2.**
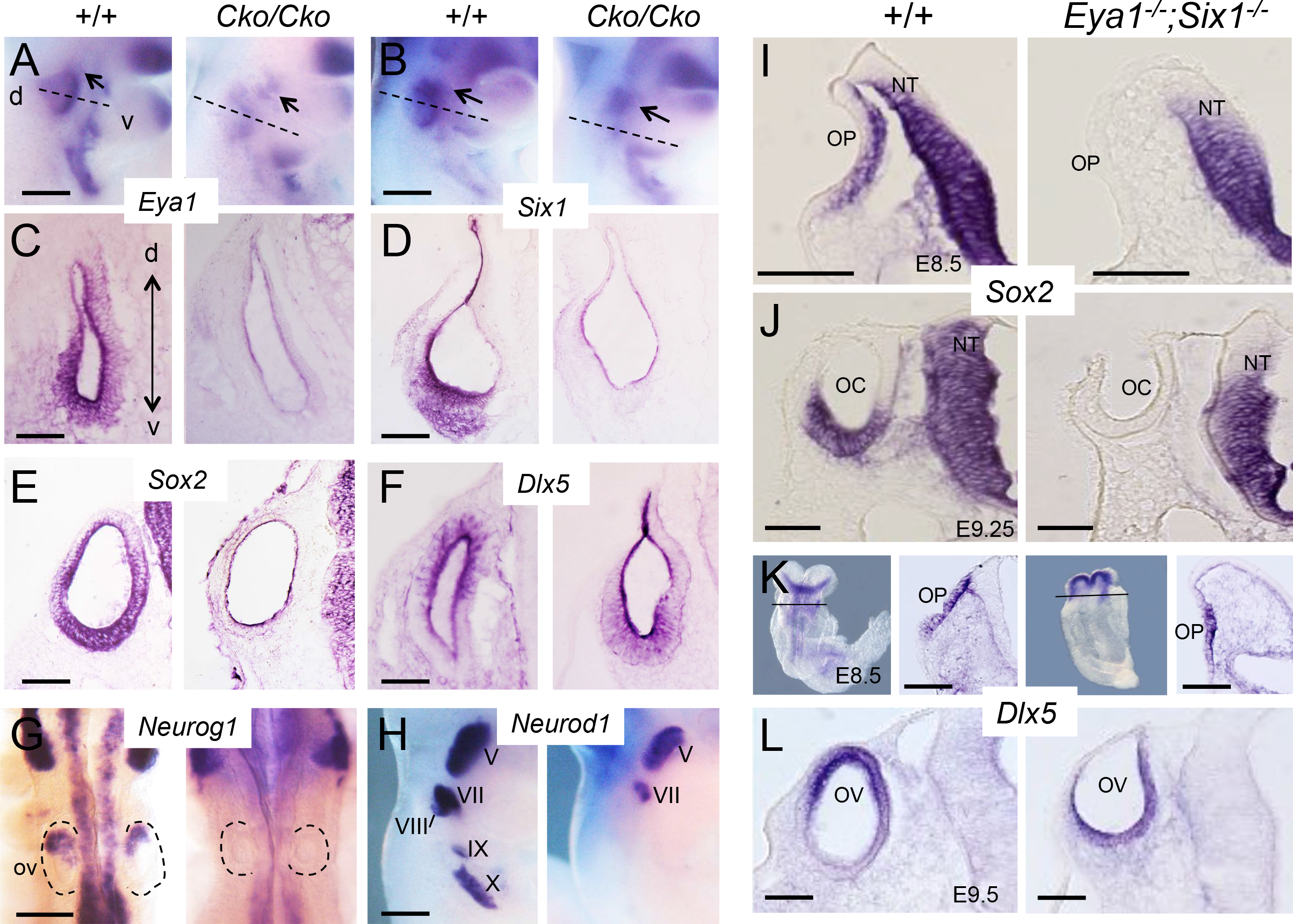
Brg1 specifies neurosensory fates in the otocyst. (A-H) Whole-mount (A,B,G,H) and section (C-F) ISH for *Eya1*, *Six1*, *Sox2*, *Dlx5*, *Neurog1* and *Neurod1* in otocyst of wild-type or *Brg1*^*Cko/Cko*^ littermates or (I-L) for *Sox2* and *Dlx5* in wild-type and *Eya1;Six1*-null embryos. Lines in K indicate regions of sections shown on the right panels. Abb.: d, dorsal; v, ventral. Scale bars: 350 μm (A,B,G,H), 50 μm (C-F, I-L).

To determine if lack of VIIIth cochleovestibular ganglion formation is due to lack of neuroblast specification or differentiation, we examined the expression of *Neurog1* and *Neurod1* by ISH. At ~E9.25, *Neurog1* expression in the anterior region of the otocyst was undetectable in *Brg1*^*Cko/Cko*^ littermates (Figure 2G), but its expression in the Vth ganglion was preserved in the CKO. *Neurod1* expression, which is detected in all cranial ganglia at E10.5 (Figure 2H), was also lost in the VIIIth ganglion of the CKO (*n*=5). These data suggest that neuroblasts are not specified in the absence of Brg1. In addition, the IXth and Xth ganglia were not formed, while the VIIth and Vth ganglia were reduced in size in the mutant (Figure 2H). Since this phenotype is similar to that seen in *Eya1*^−/−^*;Six1*^−/−^ double mutant (Zou et al., 2004), altered *Neurog1* and *Neurod1* expression could be due to loss of both *Eya1* and *Six1* expression in *Brg1*^*Cko/Cko*^. Together, these data suggest that Brg1 activity is required for the expression of *Eya1* and *Six1* to induce otic ectodermal progenitors to adopt a sensory or neuronal fate.

### *Sox2* expression depends on both Eya1/Six1 function

We have previously reported that *Sox2* expression is detectable in the otocyst of *Eya1*^−/−^ or *Six1*^−/−^ single null embryos (Zheng et al., 2003; Zou et al., 2008). Since otic neuroblast specification requires both Eya1/Six1 function, we hypothesized that Eya1/Six1 may also interact to induce *Sox2* activation in the otic placode to promote a sensory fate. Indeed, *Sox2* expression was not detected in *Eya1*^−/−^*;Six1*^−/−^ double mutant otic placode and otic cup between E8.5-9.25 (Figure 2I,J). However, *Dlx5* expression was similarly detected in the entire ectodermal region (Figure 2K,L), differing from its restricted dorsal expression in wild-type littermates (Figure 2L). In contrast, conditional deletion of *Sox2* using *Eya1*^*CreERT2*^ did not cause a ventral shift of *Dlx5* expression in the otocyst (Figure 2-figure supplement 2A-D). Both *Eya1* and *Six1* expression were observed in the ventral region of *Sox2*^*Cko/Cko*^ otocyst (Figure 2-figure supplement 2E-H), confirming that *Eya1*/*Six1* function upstream of *Sox2*. Thus, the significant reduction of *Eya1*/*Six1* expression in the *Brg1* CKO may in turn cause loss of *Sox2* expression. Together, these data indicate that Eya1/Six1 interact synergistically to induce *Sox2* expression in early otic progenitors to support the Eya1/Six1/Brg1 mediated prosensory fate specification.

### Brg1 specifies the prosensory primordium in the cochlear epithelium

We next investigated if Brg1 has a role in auditory prosensory primordum establishment during cochlear morphogenesis. The cochlear duct elongates from the ventral region of the otocyst and the prosensory progenitors proliferate until reaching a defined number. They then exit the cell cycle to become p27^Kip1+^ precursors to form a non-proliferating precursor domain in an apex to base progression between E12.5-E14.5 (Chen et al., 2002). This domain appears as a three- to four-cell layered epithelium marked by Sox2/p27^Kip1^ (Figure 3A,E). In E14.5 *Brg1* CKO given tamoxifen (Tam) at E11.75-12.5, Brg1 was selective ablation in the Eya1-expressing domains including the prosensory domain, GER and SGNs (Figure 1-figure supplement 1F,G). Sox2^+^ progenitors were present along the entire cochlear duct, which was shortened in the CKO (Figure 3B-D). However, only a few Sox2^+^ cells in the apex to medial region of the cochlea became p27^Kip1+^ (Figure 3B-D). The prosensory primordium on sections appeared slightly thinner along the basoluminal axis and Sox2 expression was maintained in the precursors in the basal layer but appeared to be downregulated in the cells in the luminal layer (Figure 3F-H), a process normally associated with HC differentiation. Within the primordial organ of Corti, only cells in the medial region were positive for p27^Kip1^ (Figure 3B,C,F-H). As cycle exit of the progenitors occurs in an apex-to-base and medial-to-lateral direction from E12.5, the few p27^Kip1+^ cells observed in the CKO cochlea could be due to the presence of Brg1 activity in those cells when CreERT2 was induced, as it takes at least 6 hrs for the CreER fusion protein to translocate to the cell nucleus and the peak marking occurs over a subsequent period of 12–24 hrs (Danielian et al., 1998). Consistent with apex to base upregulation of p27^Kip1^, we found only very few p27^Kip1+^ cells in the apical tip of the CKO cochlea after Tam injection at ~E11.75 (Figure. 3D). Thus, Brg1 is necessary to specify the prosensory primordium by regulating p27^Kip1^ activation. In the absence of Brg1, Sox2^+^ progenitors fail to become p27^Kip1+^ precursors.

**Figure 3.**
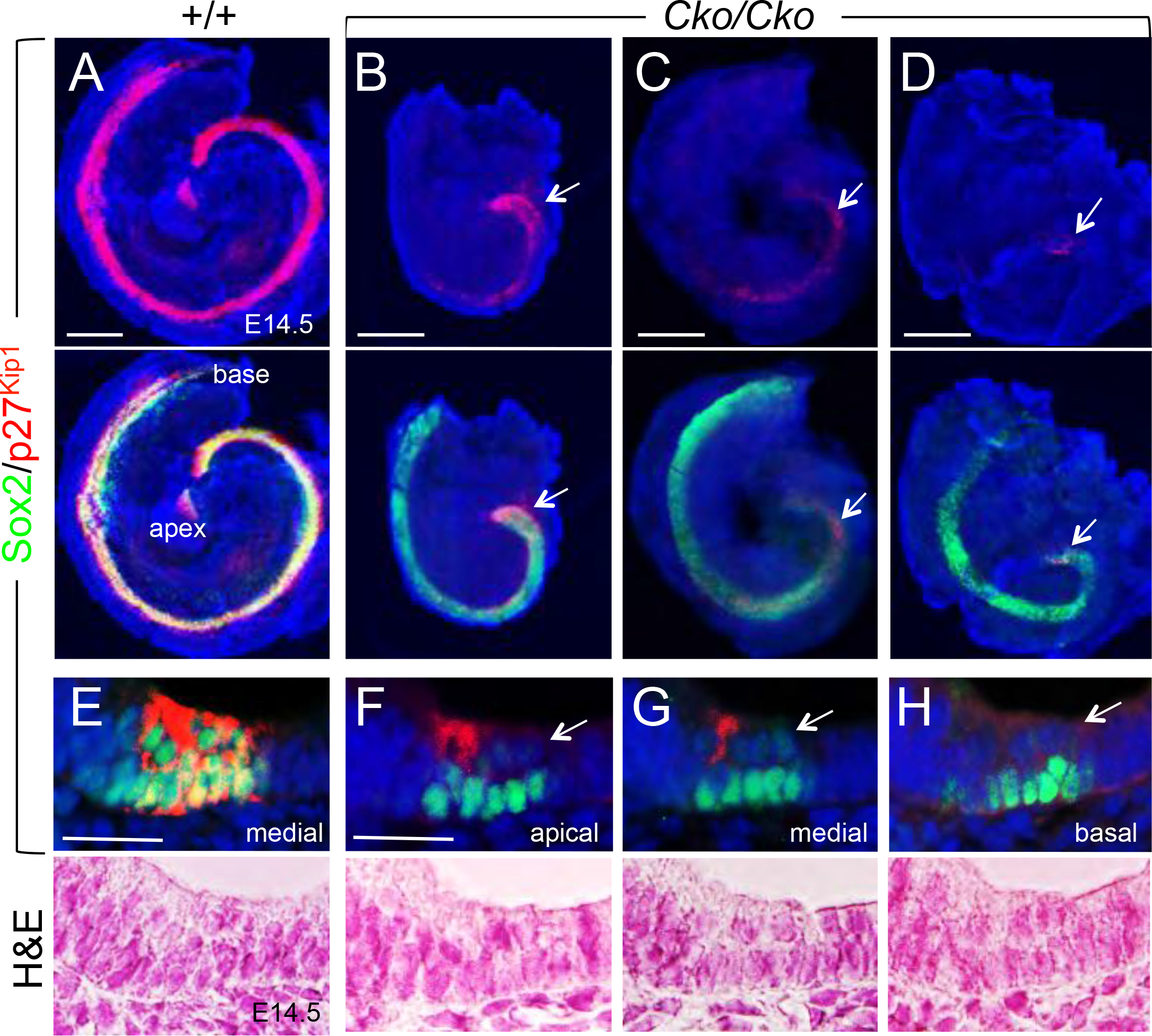
Brg1 specifies the prosensory domain in the cochlea. (A-D) Sox2/p27^Kip1^ staining of E14.5 whole-cochlea (Tam at E12.0-12.5 (A-C) or at E11.75-12.5 (D) or of cochlear sections (E-H). Lower panels of E-H are stained with H&E. Arrows point to p27^Kip1+^ cells in the apex (B-D) or reduced Sox2 in cells on the luminal surface in the sensory domain (E-H). Scale bars: 200 μm (A-D), 30 μm (E-H).

We further investigated whether the arrest of cochlear growth is due to either a decrease in proliferating cells or an increase in apoptosis. EdU incorporation demonstrated a decrease of proliferating Sox2^+^ cells in the CKO sensory epithelium at E13.5-14.5 after Tam injection at ~E11.5 (by ~83%) (Figure 3-figure supplement 3A). Quantification of the TUNEL^+^ cells in the epithelium located in the floor of cochlear duct found a strong increase in apoptosis (5.2-fold) (Figure 3-figure supplement 3B). Thus, Brg1 activity is also required for both cell proliferation and cell survival in the cochlear sensory epithelium.

### Differentiation of the postmitotic precursors into HCs or SCs is disrupted in *Brg1*^*Cko/Cko*^

We next asked whether the p27^Kip1+^ or p27^Kip1−^ precursors in the mutant presumptive organ of Corti are competent for HC or SC differentiation. Myo7a is a marker specific for differentiating HCs, while Prox1 is a SC marker specific for pillar cells lying between inner and outer HCs and three Deiters’ cells underlying outer HCs (Figure 4-figure supplement 4). Double immunostaining showed neither Myo7a^+^ HCs nor Prox1^+^ SCs present in *Brg1*^*Cko/Cko*^ cochlea at E18.5 (Tam at E11.75-12.5) (Figure 4-figure supplement 4). Sox2 expression had also disappeared in the mutant by this stage (data not shown). Thus, in the absence of Brg1 the prosensory progenitor cells fail to either exit cell cycle or enter terminal differentiation, eventually leading to loss of Sox2 expression.

**Figure 4.**
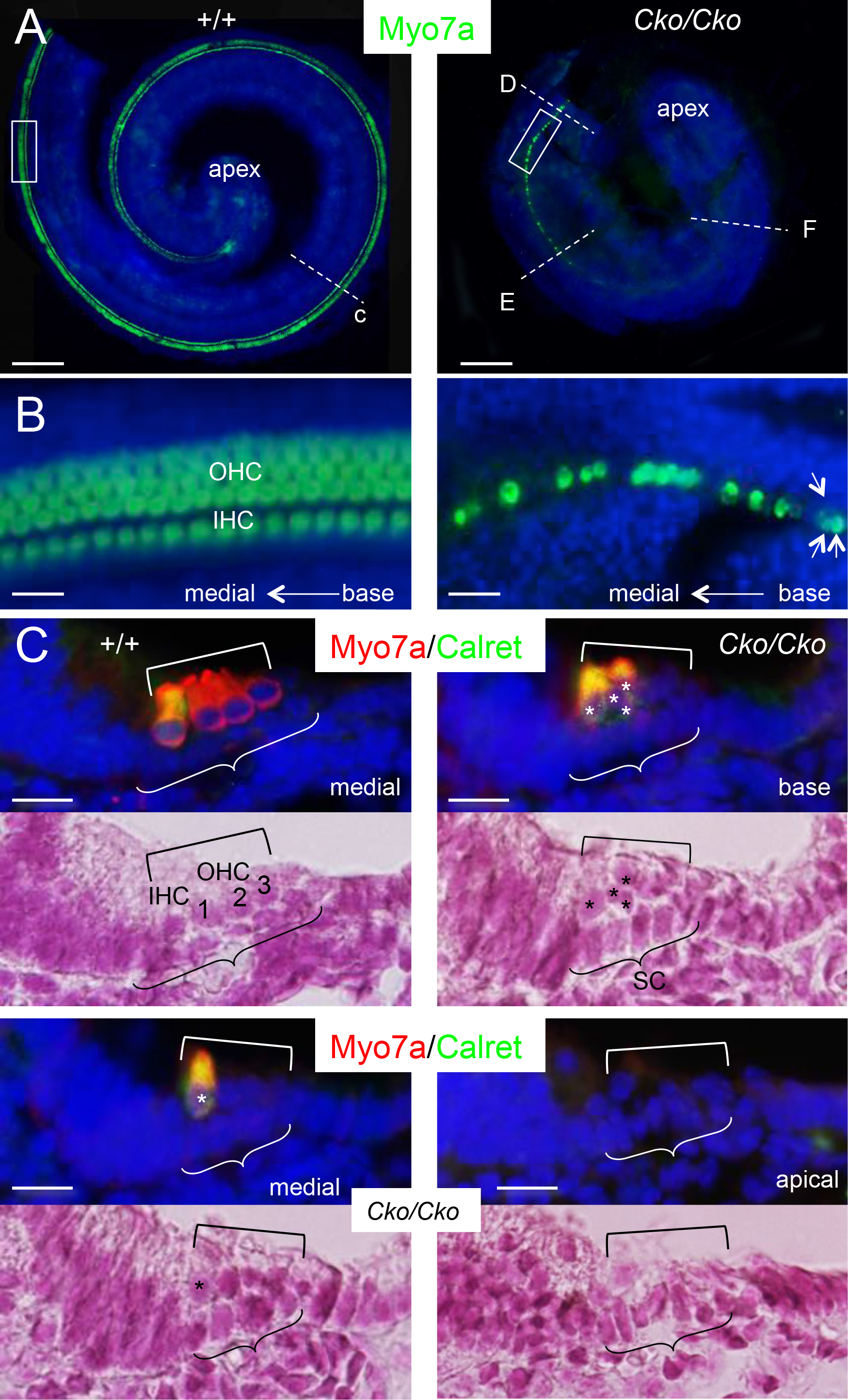
Brg1 is necessary for HC fate induction. (A) Myo7a staining of whole-cochlea at E18.0 (Tam at E13.5-14.0). (B) Higher magnification of boxed area in A. Arrows indicate disorganized Myo7a^+^ cells. (C) Myo7a/Calretinin staining on sections of E18.5 cochlea. Lower panels stained with H&E. Asterisks indicate misaligned cells in the organ of Corti. Abb.: IHC, inner HC; OHC, outer HC. Scale bars: 200 μm (A), 30 μm (B,C).

We next administered Tam more than a day later, from ~E13.5-14.0, to examine the role of Brg1 in the activation of HC and SC fates during terminal differentiation of the postmitotic precursors, which initiates near the base from ~E14.5. By E18.0, the cochlea had reached a full 1.5 turns, and differentiation of HCs, which are aligned into one row of inner and three rows of outer HCs, had reached the apical tip as outlined by Myo7a (Figure 4A,B). In *Brg1*^*Cko/Cko*^, while the cochlea was considerably smaller and only reached ~1 turn, very few HCs that failed to align into the characteristic rows of inner and outer HCs were present in the base, and none were observed in the medial to apex (Figure 4A,B). Since HC differentiation occurs in a medial-to-lateral (inner-to-outer) gradient, the Myo7a^+^ HCs likely represent the inner HCs, which begin to differentiate at least more than a day prior to outer HCs. Indeed, immunostaining for calretinin, a marker specific for inner HCs, confirmed that all Myo7a^+^ HCs in the base were also calretinin^+^ (Figure 4C). As seen on sections, the organ of Corti was patterned into a two-cell layered epithelium with one-cell layer of HCs on the luminal side and one cell layer of underlying SCs (Figure 4C). In *Brg1*^*Cko/Cko*^, no obvious change in the epithelial thickness along the basoluminal axis was observed. However, cells in the presumptive organ of Corti were irregularly patterned and disorganized with morphological alteration in the CKO compared to those in the control littermates (Figure 4C). Thus, Brg1 is necessary for HC fate induction and sensory epithelium formation in the cochlea.

The SCs maintain high levels of p27^Kip1^ and Sox2 through adult stage (Figures 5A,B). Although the SCs in the CKO were irregularly shaped and the organ of Corti became slightly narrower compared to that in wild-type controls, they expressed Sox2/p27^Kip1^. Prox1 was expressed in *Brg1* CKO SCs in the base but not in the medial to apex, and the Prox1^+^ cells were irregularly aligned (Figure 5C). However, the neurotrophin receptor P75^NTR^ expression in inner pillar cells was not affected in the CKO (Figure 4D). The glutamate-aspartate transporter (GLAST) expression in inner border and inner phalangeal cells were detectable in the CKO cochlea, but they appeared to be disorganized and dislocated on the luminal layer (Figure 4E). The calcium-binding protein S100A marks inner HCs and surrounding inner border, inner phalangeal and inner pillar cells, as well as the Deiters’ cells underlying the outer HCs (Figure 5E,F). S100A^+^ cells were present in the CKO, but they were disorganized with more than one cell on the luminal side. The S100A^+^ cells on the luminal side where outer HCs are normally located were also Myo7a^+^ cells in the base, but not in medial cochlear duct (Figure 5F). From medial to apical end, all S100A^+^ cells on the luminal layer were Myo7a^−^ cells. In agreement with the calretinin staining (Figure 4C), the S100A^+^ cells on the luminal layer likely represent the inner HC precursors that failed to differentiate into Myo7a^+^ cells. Nonetheless, our data indicate that Sox2^+^p27^Kip1+^ SCs in the *Brg1*^*Cko/Cko*^ failed to fully differentiate into Prox1^+^ SCs, resulting in disorganized sensory epithelium.

**Figure 5.**
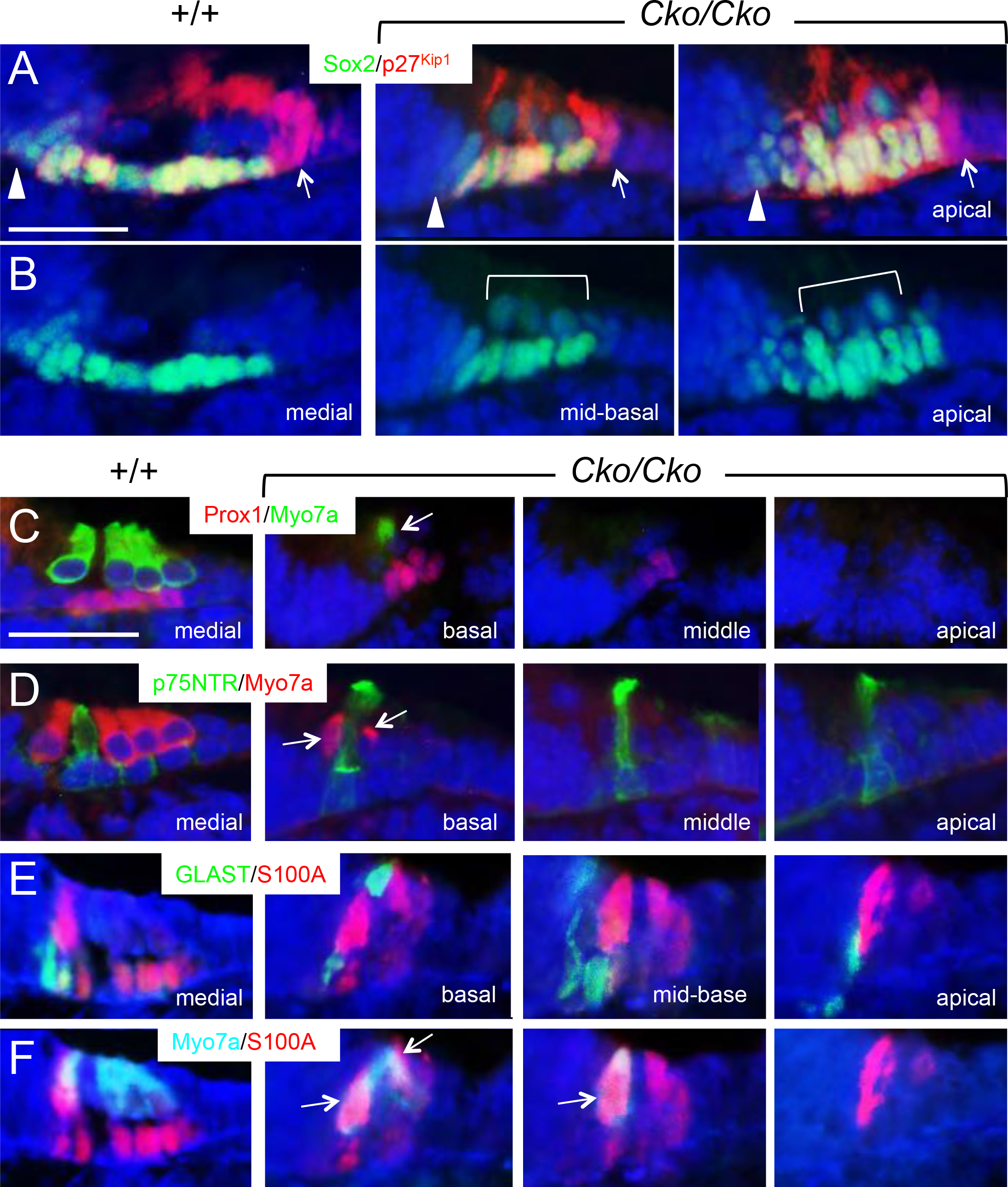
Differentiation of supporting cells is disrupted in the absence of Brg1. Antibody labeling for p27^Kip1^ (red) and Sox2 (green) (A,B), Prox1 (red) and Myo7a (green) (C), p75^NTR^ (green) and Myo7a (red) (D), GLAST (green) and S100A (red) (E), Myo7a (cyan) and S100A (red) (F) in the organ of Corti in wild-type and *Brg1* CKO (*Eya1*^*CreER*^*;Brg1*^*fl/fl*^) littermates given tamoxifen from E13.5-14.0. Arrowheads in a point to Sox2^+^ but p27^Kip1−^ cells in the GER, while arrows point to p27^Kip1+^ but Sox2^−^ Hensen’s cells. In the mutant, the organ of Corti appears narrower. Arrows in panel D point to cells positive for Myo7a and in G point to cells positive for both Myo7a and S100A. Scale bars: 30 μm.

In contrast to Sox2 expression in the CKO cochlear sensory epithelium (Figure 5A,B), the expression of *Eya1*, which is normally expressed in differentiating hair cells at E17.5-18.5 (Zou et al., 2008), was markedly reduced and only some faint signal was observed in the basal region (arrow, Figure 6A). *Six1* is expressed in both hair and supporting cells (Zhang et al., 2017), but its expression in both cell types was almost undetectable in the CKO cochlea (Figure 6A). In addition, *Atoh1* expression was undetectable in the CKO (Figure 6A). This indicates that in the absence of Eya1/Six1, Sox2 alone is insufficient to induce *Atoh1* expression in Sox2^+^ progenitors to promote a HC fate in vivo.

**Figure 6.**
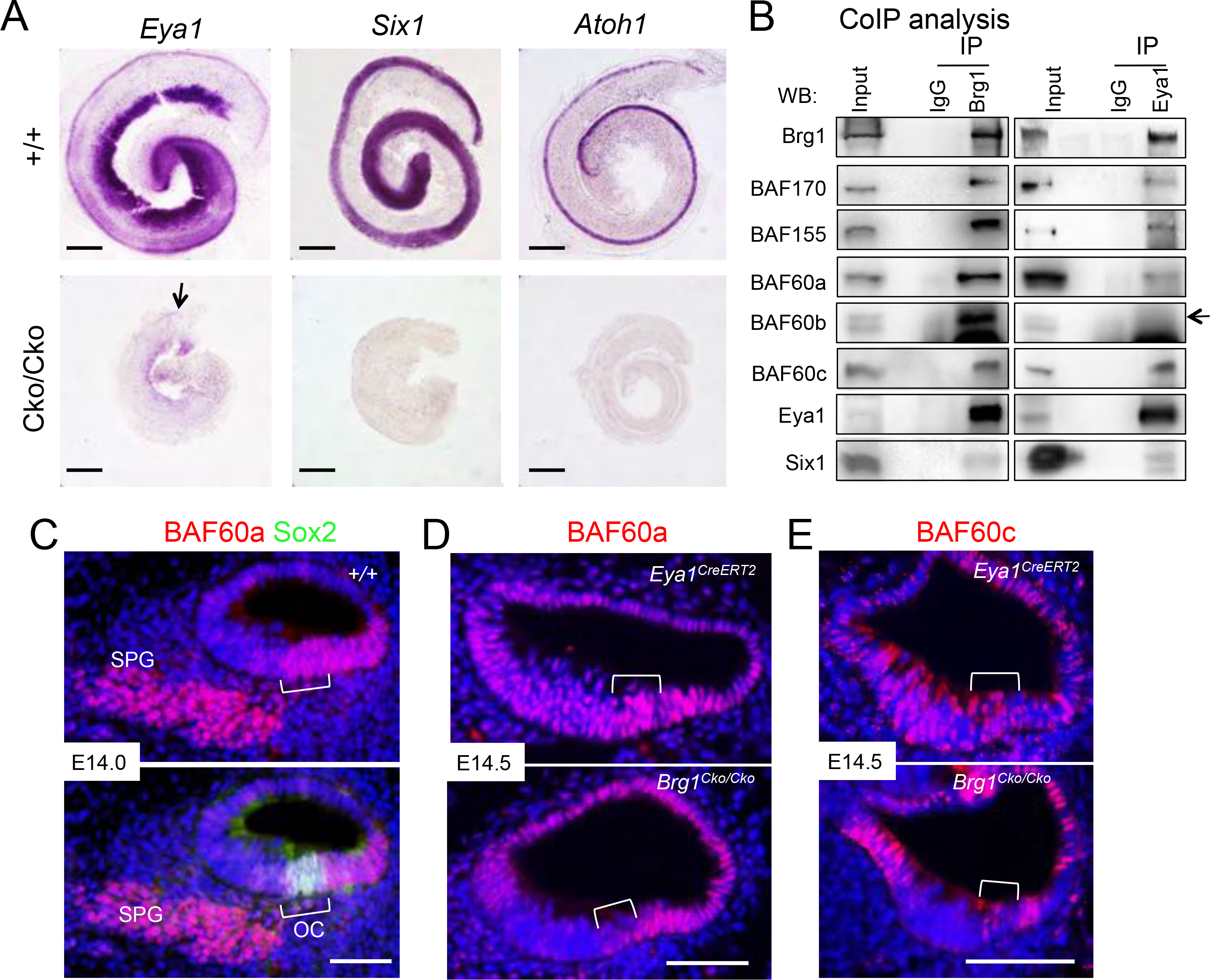
Brg1 is necessary for maintenance of *Eya1*, *Six1* and *Atoh1* expression in the cochlear sensory epithelium and Eya1 selectively interacts with BAF60a/c but not BAF60b. (A) In situ hybridization for *Eya1*, *Six1* and *Atoh1* in cochlea of wild-type or *Brg1*^*Cko/Cko*^ littermates at E17.5 given tamoxifen from E13.5. (B) CoIP analysis. Nuclear extracts were prepared from E13.5 cochleae. Antibodies for IP and western blot are indicated. (C,D) Immunostaining for(C) BAF60a/Sox2 or (D) BAF60a alone on sections of wild-type cochlea at E14.0, *Eya1*^*CreERT2*^ or *Brg1*^*Cko/Cko*^ (*Eya1*^*CreERT2*^*;Brg1*^*fl/fl*^) embryo at E14.5 given tamoxifen at E11.5-12.5. (E) Immunostaining for BAF60c in control (*Eya1*^*CreERT2*^) or *Brg1*^*Cko/Cko*^ littermates given tamoxifen at E11.5-12.5. Scale bars: 200 μm (A), 50 μm (C-E).

### Eya1-Six1 simultaneously interacts with the BAF60a/c structural subunit

The SWI/SNF complexes contain more than 10 associated BAF factors, which generate a variety of subcomplexes involved in dynamic and tissue-specific interactions with TFs and other chromatin-modifying enzymes. Some BAFs have variants that are selectively used in cell fate commitment (Ho and Crabtree, 2010). Peptides from BAF170, BAF155 and BAF60b were detected in our proteomics analysis of the Eya1 complexes purified from HEK293 cells that constitutively express a multi-tagged-Eya1 (2×HA-3×Flag-Eya1) (J. Xu et al., manuscript in preparation), leading to the hypothesis that Eya1 activity may depend on the activity of such BAFs. We therefore examined whether Eya1 interacts with such BAFs in the cochlea. We have generated the 2×HA-3×Flag-Eya1 knockin mouse line (J. Li et al., manuscript in preparation), which allows us to use anti–Flag or–HA for immunoprecipitating the multi-tagged Eya1 protein complexes. CoIP analyses using nuclear extracts from E13.5 cochleae confirmed physical association of Eya1 with Brg1, BAF170 and BAF155 (Figure 6B). BAF60 has three variants, a, b and c, and in contrast to our mass spectrometry data showing interaction of Eya1 with BAF60b in 293 cells, Eya1 associated simultaneously with BAF60a/c but not BAF60b in the cochlea (Figure 6B). Consistent with our previous observation (Buller et al., 2001), Eya1 formed a complex with Six1, which showed similar physical interaction with BAF60a/c but not with BAF60b (Figure 6-figure supplement 5A). In contrast, Brg1 physically associated with all three BAF60 variants along with BAF170/BAF155 (Figure 6B). This suggests formation of specific BAF60a/c-based SWI/SNF-Eya1/Six1 complexes.

We next investigated their expression pattern during cochlea development. BAF60a showed higher levels of expression in the prosensory primordium (Figure 6C,D) and later in both HCs and SCs (Figure 6-figure supplement 5B), while BAF60c appeared to be expressed in discrete cells in the prosensory domain (Figure 6E) and by E18.5 its expression was only detected in the HCs and pillar cells Figure 6-figure supplement 5C). Both BAF60a/c are also expressed in Hensen’s cells flanking the outer HCs and in SGNs (Figure 6-figure supplement 5B,C). In *Brg1*^*Cko/Cko*^ cochlea at E14.0-14.5 (Tam at E11.5), both BAF60a/c levels were reduced specifically in the *Eya1*-expressing domain (GER and prosensory primordium) (Figure 6D,E). However, in contrast to marked depletion of Brg1 in the prosensory progenitors (Figure 1-figure supplement 1G), some BAF60a^+^ cells and clusters of BAF60c^+^ discrete cells were detectable in the CKO prosensory primordium. While it is possible that other ATPase-based SWI/SNF complexes containing BAF60a/c may form in different subsets of the progenitors, our data suggest that in the absence of Brg1-BAF complex assembly, BAF60a/c may become unstable and undergo degradation.

Both BAF170/BAF155 were also detected in the prosensory primordium (Figure 6-figure supplement 5D,E). By E18.5, high levels of BAF170 were maintained in HCs and SCs and in GER as well as in stria vascularis and SGNs (Figure 6-figure supplement 5F). In contrast, BAF155 was weakly expressed in HCs and SCs but strongly in Henen’s cells and LER (Figure 6-figure supplement 5G). Together, these data provide insight into how distinct BAF subunits might be used to generate a diversity of SWI/SNF complexes during cochlea morphogenesis to promote lineage-specific properties.

### Role for Brg1 in global gene activation in the cochlea

To explore Brg1 genome-wide binding profile and its association with regulatory sequences in the prosensory progenitors, we performed ChIP-seq on E13.5 cochleae. MACS (Zhang et al., 2008) peak calling identified 8086 (with a *P* value threshold of 1e-5) and 1292 (a *P* value threshold of 1e-6) Brg1 peaks (Figure 7A). ~28% of Brg1-bound regions were within 5 kb of promoter-5’ UTR and ~72% were located within 5-500 kb of TSSs of the nearest gene (Figure 7B). Analysis of the 8086 Brg1-enriched regions found 3079 (~28% promoter-proximal and ~72% distal) associated with H3K27ac, a well characterized epigenetic mark associated with transcriptionally active chromatin (Figures 7A and Figure 7-figure supplement 6A). Consistent with Brg1 involvement in active and repressive gene expression in other tissues (Attanasio et al., 2014; Trotter and Archer, 2008), our cochlear ChIP-seq data indicate that Brg1 is associated not only with transcriptionally active chromatin, but also with other regulatory regions.

**Figure 7.**
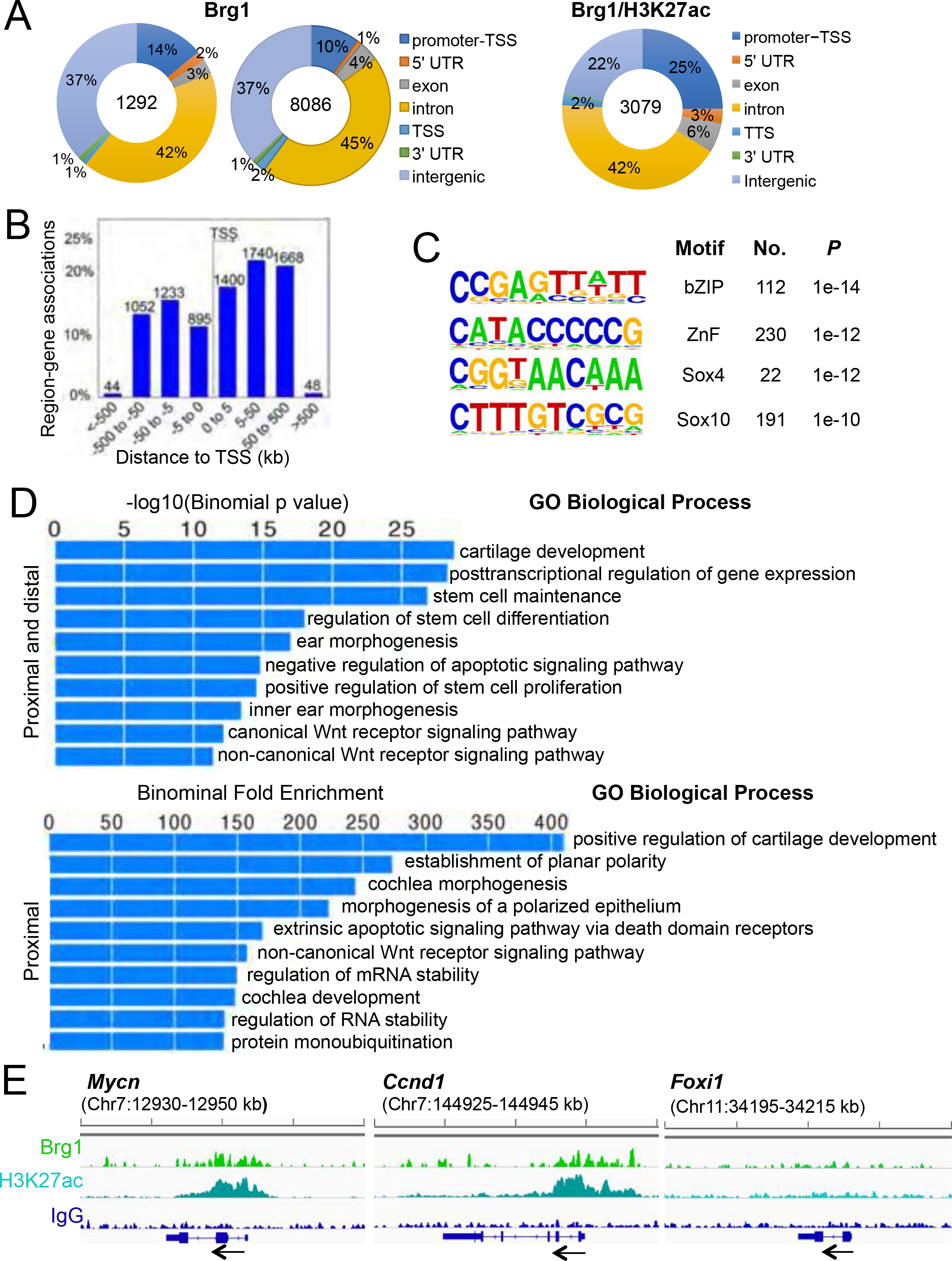
Genomic binding features of Brg1 in E13.5 cochlea. (A) Genomic distribution of Brg1-enriched regions and its overlap peaks with H3K27ac. (B) Distribution of Brg1 peaks relative to TSSs. (C) The top three motifs identified by Homer tools. (D) Differential enrichment for functional annotation terms associated Brg1-enriched regions. Shown are the top 10 enriched “Biological process terms” associated with the 1292 Brg1 enriched regions or specific to proximal Brg1 regions within 1 kb from promoters. (E) Genomic view of Brg1 enrichment at *Mycn* and *Cyclin D1* loci but not at *Foxi1* locus.

To explore whether Brg1-enriched sites contain consensus motifs and to reduce the number of regions for motif analysis, we examined the 1292 peaks for the presence of known motifs. The top three motifs contain consensus sequences for bZip proteins, Zinc finger proteins and SOX factors (Figure 7C), all of which are known to specifically interact with Brg1-containing SWI/SNF and act as Brg1’s transcriptional coregulators (King and Klose, 2017; Marathe et al., 2017; Trotter and Archer, 2008). Thus, this analysis provides insight into potential DNA-binding TFs that Brg1 interacts with at gene promoters and distal regulatory elements in the cochlea.

Gene Ontology (GO) analysis showed highly enriched “biological process terms” near genes associated with processes involving development and morphogenesis, including inner ear and cochlea specific processes characteristic of cochlea development (Figure 7D and Figure 7-figure supplement 6C). Brg1 binding was evident at genes involved in cell division such as *Mycn* (*Nmyc*) (Figure 7E) and *Myc* (*cMyc*) (Figure 7-figure supplement 6D) as well as at their downstream targets *Cyclin D1*/*D2* (Figures 7E and Figure 7-figure supplement 6D). Genes related to inner ear/cochlea morphogenesis and development include *p27*^*Kip1*^, TFs (such as *Eya1/4, Six1/4, Sox2/4/5/9, Gata3*, *Sobp, Prox1, Id2/3*, and *Tbx18*), and genes participating in Wnt signaling (*Wnt5a/Fzd/Ptk7/Sfrp2/Cthrc1/Ptk7)*, Hh *(Gli3/Ptch1)*, Fgf-signaling (*Fgfr1/2/Fgf9*), and Notch pathway (*Jag1/Lfng/Dll1/Hes1/Hey1/Hey2/Notch1/2*) (Supplementary files 1-3 and Figure 7-figure supplement 7). In contrast, no Brg1 peaks were identified at genes that are not expressed in the cochlea such as at *Foxi1* (Figure 7E) and *Six6* (Figure 7-figure supplement 6D) loci. Together, our ChIP-seq analysis not only indicates Brg1-binding of key neurosensory genes including *Eya1*, *Six1*, and *Sox2* but also uncovers a robust set of Brg1-bound regions near genes involved in expected tissue-specific biological processes.

### ChIP assays confirm Brg1 binding at discrete regulatory regions in the *Eya1*, *Six1*, *Sox2*, *Eya4*and *p27^Kip1^* loci and co-occupancy by BAF60a/c

At the *Eya1* locus, Brg1-bound sites were recovered in promoter (I) and intergenic (II) regions and most of these regions are ECRs (evolutionary conserved regions) based on comparison of mouse with human genomes (Figure 8A). Since *Eya1* expression in the otocyst also requires Brg1 activity, we speculated that Brg1 may bind to the same elements at the *Eya1* locus to activate its expression in early otic progenitors and to maintain its expression in cochlear sensory epithelium. ChIP-qPCR assays on otocysts dissected from E10.5 embryos confirmed strong Brg1 enrichment at the promoter (+610 to +429 bp; +2 to −180 bp; −373 to −532 bp) (primer positions, Supplementary file 4) in wild-type samples (Figure 8B). For the intergenic sites, ChIP-qPCR revealed strong association of Brg1 with an ECR (−913924 to −914124 bp), but very weak enrichment at two non-ECRs (−196849 to −197024 bp and −197523 to −197695 bp) that had relatively weaker Brg1 signals in ChIP-seq (red stars). ChIP-qPCR on *Eya1*^*CreERT2*^*; Brg1*^*fl/fl*^ littermates showed a significant reduction in Brg1-enrichments at these regions (Figure 8B). Similarly, ChIP-qPCR for four regions at the *Six1* locus confirmed Brg1 binding to two of the downstream intergenic elements (+12227 and +8299) and disruption of Brg1 association with these regions in *Brg1* CKO otocysts (Figure 8-figure supplement 8A,D). These results further support that Brg1-occupancy is required for the activity of these genes during inner ear development.

**Figure 8.**
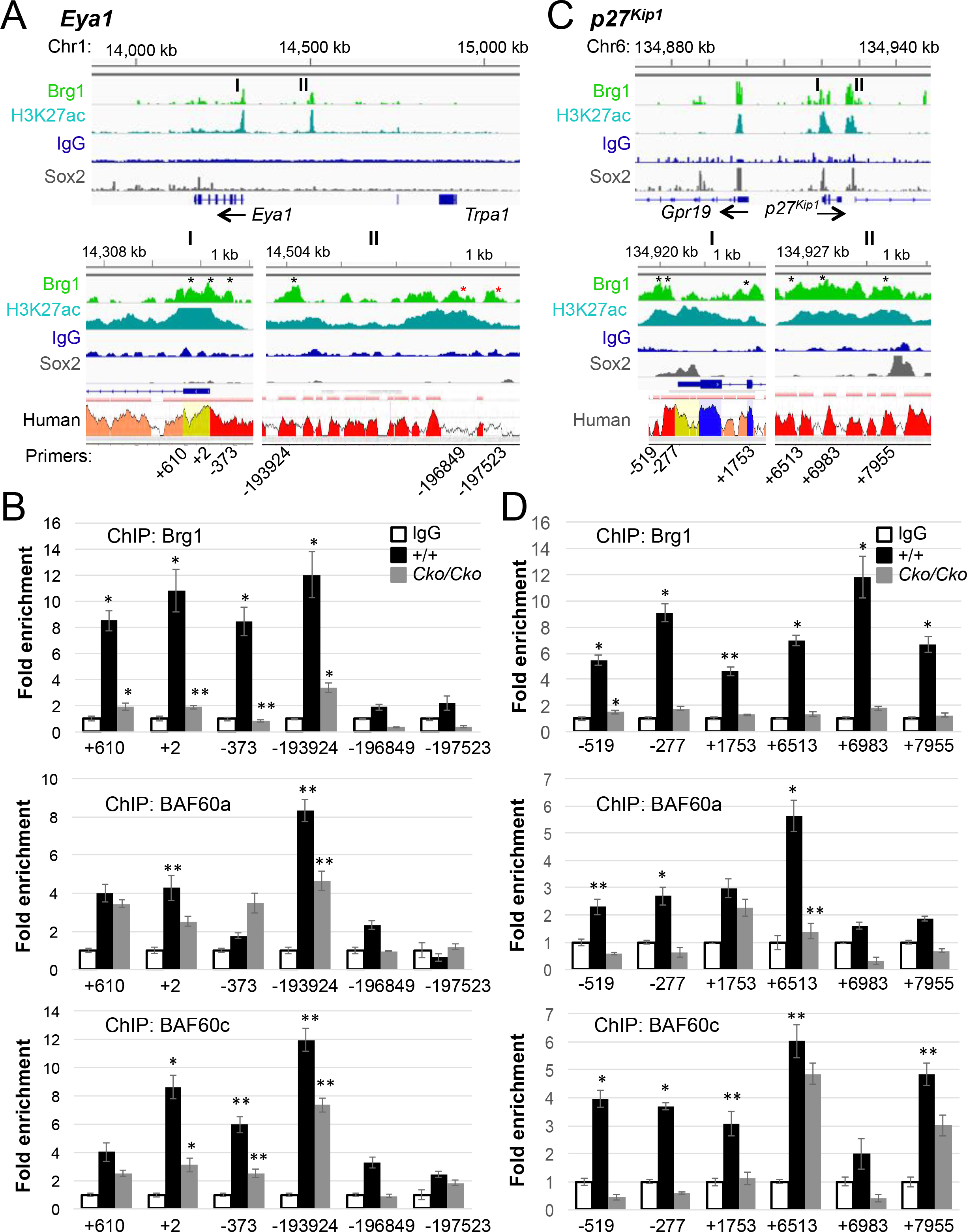
Brg1-bound regions at the *Eya1* and *p27Kip1* loci and co-occupancy patterns of BAF60a/c at these sites. (A,C) Brg1-bound regions at promoter-proximal (I) and distal (II) regions at the *Eya1* (A) or *p27*^*Kip1*^ (C) loci and alignment of these regions with published Sox2 ChIP-seq (Kwan et al., 2015) and from mouse and human. Peak heights indicate degree of sequence homology; pink bars above the peaks denote ECRs; UTR (yellow) correspond thin blue box in IGV view; exons (thick blue boxes). Peak regions corresponding to ECRs (black asterisks) or non-ECRs (red asterisks) were selected for ChIP analysis. See Table S4 for positions of these sites. (B,D) ChIP-qPCR data using chromatin derived from otocysts of wild-type or *Brg1*^*Cko/Cko*^ littermates at E10.5 (Tam at E8.0-9.0) (B) or from cochleae of wild-type or *Brg1*^*Cko/Cko*^ littermates at E14.5 (Tam at E11.5-12.5) (D) and anti-Brg1, anti-BAF60a or anti-BAF60c antibodies. Data were normalized with inputs and mock IgG control. Note very faint or no enrichment of Brg1 at the non-ECRs of *Eya1*, and weak enrichment for BAF60a at −373 bp of the *Eya1* and at +6983 bp and +7955 bp of the *p27*^*Kip1*^ and for BAF60c at +6983 bp of the *p27*^*Kip1*^. Representative of three independent experiments, n=3 for each group, \**P*<0.01, \*\**P*<0.05 determined by Student’s t-test).

At the *Sox2* locus, Brg1-binding was recovered at an intergenic region located at ~691 kb upstream from the promoter (Figure 8-figure supplement 8B). This region was also recovered in the Brg1 ChIP-seq data on E11.5 forebrain (Attanasio et al., 2014). ChIP-qPCR within 1 kb of the Brg1-bound region (asterisks, Figure 8-figure supplement 8B) confirmed strong Brg1 binding at +691335 and +191656 in wild-type otocyts and large reduction of Brg1 binding at these sites in *Brg1* CKO (Figure 8-figure supplement 8E), thus suggesting possible requirement of Brg1-occupancy at this region for *Sox2* expression in the otic progenitors. ChIP assay also confirmed Brg1-binding at the *Eya4* gene (Figure 8-figure supplement 8C,F), which is known to have a role in sensory cell development as *EYA4*-deficiency in humans causes hearing impairment (Pfister et al., 2002).

To characterize if BAF60a or BAF60c is a component of the SWI/SNF complex binding at the *Eya1* locus, we performed ChIP assay with anti-BAF60a or BAF60c respectively. BAF60a showed association to other three Brg1-bound sites (+610, +2 and −193924; Figure 8B), while BAF60c revealed enrichment at all four sites with different affinity levels. *Brg1*-deficiency led to ~2 to 4-fold reduction in BAF60a or BAF60c enrichment at these sites. The co-occupancy patterns of BAF60a/c on Brg1-bound sites could result from transient binding by separate SWI/SNF components in different subsets of Eya1^+^ cells. In summary, the ChIP experiments on E10.5 otocysts confirm enrichment of Brg1 at the discrete ECR regions in the *Eya1* locus from early stage. Furthermore, our results support that Brg1 is the preferred ATPase subunit of the SWI/SNF complex binding at the discrete regulatory regions of *Eya1* to regulate its expression.

The cyclin-dependent kinase inhibitor p27^Kip1^ (Cdkn1b) is required to correctly time cell cycle exit of the prosensory progenitors prior to their differentiation and is regulated by Sox2 in cochlear SCs (Liu et al., 2012). Brg1 ChIP-seq recovered strong peaks at the discrete promoter-proximal and distal regions in the *p27kip1* locus (Figure 8C). We compared these regions with public Sox2 ChIP-seq data from immortalized Sox2^+^ otic progenitor cells (Kwan et al., 2015) and found that Sox2-bound sites at the promoter and distal sites overlap with the Brg1-bound regions (Figure 8C), suggesting that Sox2 may interact with Brg1-BAF complexes to regulate *p27*^*Kip1*^ expression. Indeed, Sox2 physically interacts with Brg1 in the cochlea by coIP analysis (Figure 6-figure supplement 5A). ChIP assays on E13.5 cochleae confirmed association of Brg1 to the promoter-proximal and distal sites (−519, −277, +1753, +6513, +6983 and +7955) (Figure 8C,D). *Brg1*-deficiency (given Tam from E11.5) significantly decreased Brg1 binding at these sites. Thus, reduced p27^Kip1^ expression in *Brg1* CKO cochlea (Figure 3) is strongly associated with decreased Brg1 binding to the discrete regulatory regions of *p27*^*Kip1*^ gene. *Brg1*-deficincy also led to decreased BAF60a or BAF60c enrichment at these Brg1-bound sites. Together, these co-occupancy patterns at the ECRs further support that Brg1-based SWI/SNF complex containing BAF60a or BAF60c is necessary for *p27*^*Kip1*^ expression through binding to the discrete regulatory regions in the *p27Kip1* locus.

## Discussion

We have previously shown that Eya1-Six1 interacts with Brg1-BAFs to convert cochlear nonsensory GER cells to SGNs, yet nothing is known about the in vivo requirement of the Brg1-BAF complexes for inner ear neurosensory cell fate commitment and their involvement in regulatory pathways controlling inner ear formation. Here we demonstrate that otic ectoderm requires the chromatin-remodeling enzyme Brg1 to form inner ear structures. The pattern of cochlear Brg1 genomic binding and its strong association with H3K27ac suggest that Brg1 is substantially involved in regulatory pathways governing inner ear development.

Ablation of Brg1 in *Eya1*-expressing cells blocked the activation of neurosensory cell fate inducing programs. This phenotype is linked to loss of *Eya1*/*Six1* expression, which results in loss of *Sox2* and *Neurog1* expression. Eya1, Six1 and Sox2 form the core transcription circuitry for neurosensory precursor cell fate specification and subsequent differentiation in the inner ear. However, to date, nothing is known about what lies upstream of these genes in the inner ear. Our findings provide the first in vivo evidence that Brg1-based SWI/SNF chromatin remodeling factors regulate the expression of *Eya1*, *Six1* and *Sox2* at different stages of inner ear development. Our data show that Brg1 plays a crucial role in coordinating multiple processes during otic neurosensory cell specification and differentiation.

How do the Brg1-based SWI/SNF complexes help to establish and maintain neurosensory cell identities? SWI/SNF chromatin remodeling complexes are typically recruited to genomic sites via physical interactions with sequence-specific TFs to execute spatiotemporal control over specific target genes during developmental processes (Ho and Crabtree, 2010; Neely et al., 1999). Therefore, Brg1 may differentially impact the spatial and temporal expression of Eya1/Six1 in the otic ectoderm. The otic placode forms from a common pre-placodal ectoderm, which is defined by the members of Eya and Six families. However, unlike *Six1/4* and *Eya2* that are coexpressed in the pre-placodal ectoderm, the mouse *Eya1* is expressed in the otic and epibranchial placodal ectodermal regions (Chen et al., 2009). Thus, Eya1 may have a specific role in defining an otic fate from the pre-placodal ectoderm and the timing of Eya1 activation may require a unique Brg1-SWI/SNF complex in response to external signals. In support of this, ChIP assays on E10.5 otocysts confirmed Brg1 association with the promoter and distal regulatory regions at the *Eya1* locus. Future work is required to reveal exactly how the binding of Brg1 to *Eya1* locus is related to establishing otic fate. This process likely involves other TFs that are expressed in the ectoderm, during the transition from the pre-placodal stage to Eya1^+^ multipotent otic progenitors before neurosensory cell fate induction.

Once Eya1 is activated, Brg1-BAFs may alter the structure of chromatin at lineage-specific genes to facilitate the function of Eya1, which interacts with Six1 and other factors. Our observation of a direct physical interaction between Brg1-BAFs and Eya1-Six1 suggests that Eya1-Six1 use the chromatin-remodeling properties of Brg1-BAF complexes to regulate transition from a multipotent otic ectodermal cell to a prosensory restricted progenitor cell by activating lineage-specific genes such as Sox2 or to a committed neuronal fate by activating Neurog1-Neurod1. This explains why ablation of both Eya1/Six1 also leads to a lack of *Sox2* (Figure 2) or *Neurog1-Neurod1* (Ahmed et al., 2012b) expression. Since the expression of *Eya1*/*Six1* is markedly reduced in *Brg1* CKO, lack of Brg1-SWI/SNF binding at these genes could prevent their activation. This in turn simultaneously prevents expression of *Sox2* and *Neurog1-Neurod1* and results in a developmental block of neurosensory cell fate specification. We thus speculate that Brg1 may bind to the regulatory elements at the *Eya1* and *Six1* loci in order to activate their coexpression before neurosensory fate induction in otic placode and to maintain the coexpression of these genes in otic ectoderm and cochlear sensory epithelium at later stages. Future studies using a combination of genomic and molecular approaches are required to reveal the molecular details of TFs and Brg1-BAF complexes involved in activating *Eya1* or *Six1* during early otic fate induction.

Our data also show an essential role for Brg1 in both cell survival and proliferation within the otic ectoderm, which is consistent with Brg1’s necessity in cell survival and proliferation in various tissues. As Eya1 or Six1 is also required for normal cell proliferation and survival in the otic ectoderm (Zheng et al., 2003; Zou et al., 2006), Brg1 may cooperate with Eya1/Six1 to regulate these processes, which explains why loss of any of these genes causes a decrease in cell number in the otic ectoderm and leads to growth arrest. Ablation of Brg1 in the developing cochlea at later stages underscores the continuous contribution of Brg1-BAF remodelers during prosensory precursor cell differentiation, from the regulation of precursor proliferation and survival to the induction of cell type-specific genes.

In gain-of-function studies, we previously demonstrated the requirement of Brg1 activity and its interaction with Eya1/Six1 in inducing a robust neuronal differentiation program from cochlear nonsensory GER cells or ectodermal cells surrounding the otocyst in explants (Ahmed et al., 2012b). However, it remained unknown whether Brg1 is also necessary for HC fate induction. The present loss-of-function study now demonstrates the necessity of Brg1 in inducing auditory sensory epithelium formation. Brg1 is not only essential for specification of the prosensory domain but also for terminal differentiation of precursor cells within the domain into either HCs or SCs. Conditional deletion of Brg1 between E11.75−12.5 prevented Sox2^+^ prosensory progenitors from becoming p27^Kip1+^ postmitotic precursors. Thus, Brg1-BAF complexes may facilitate regulation of *p27*^*Kip1*^ expression and other downstream genes by Sox2, thus explaining why Sox2 alone is insufficient to induce p27^Kip1^ expression. This is consistent with the physical interaction between Sox2 and Brg1 and Sox2 co-occupancy at a subset of Brg1-bound regions at the *p27*^*Kip1*^ locus. The necessity of Brg1 activity in *p27Kip1* regulation is further supported by the marked reduction in Brg1 enrichment at discrete regulatory ECRs in the *p27Kip1* locus in the *Brg1* CKO cochlea (Figure 8D). Interestingly, when Brg1 was deleted at later stages between E13.5-14.0 after *p27*^*kip1*^ activation, p27^Kip1+^ cells also failed to fully differentiate into Myo7a^+^ HCs or Prox1^+^ SCs. Although this developmental defect may have resulted from a failure to appropriately regulate chromatin structure, it may also be attributed to a deficiency in modification of other proteins that are crucial for HC fate induction or SC differentiation, such as Eya1, Six1, Atoh1 and Prox1, all of which are not expressed in the *Brg1* mutant. In the absence of Eya1 and Six1 in *Brg1* CKO cochlea, Sox2 alone appears to be incapable of activating Atoh1 or maintaining Prox1 expression in Sox2^+^ precursors to promote HC fate induction or SC differentiation. Although it is currently unclear whether Eya1-Six1 also use Brg1 activity to facilitate *Atoh1* activation and HC induction during cochlea development, in our ChIP-seq data sets, weak enrichment of Brg1 and H3K27ac was observed at the 1.4 kb *Atoh1* HC-specific enhancer located ~3.4 kb 3’ of the *Atoh1* coding sequence (red star, Figure S8). This enhancer mediates transcriptional regulation by Eya1-Six1-Sox2 (Ahmed et al., 2012a). It is possible that *Atoh1* may not be activated yet in the cochlear samples collected from E13.5 embryos for our ChIP-seq, as suggested by the weak H3K27ac signals at this enhancer. This is consistent with the observation that *Atoh1* activation is induced between E13.5-14.5 (Chen et al., 2002). We are currently performing Brg1 ChIP-seq on cochleae at later stages between E14.5 to E15.5 to examine if Brg1 enrichment at this HC-specific *Atoh1* enhancer increases. Stronger Brg1 binding at later stages would suggest that recruitment of Brg1-BAF complex to the 1.4 kb HC-specific enhancer may confer temporal control of HC fate activation by Eya1-Six1-Sox2. Nonetheless, stronger Brg1 enrichment at the promoter-proximal regions of the *Atoh1* suggests that Brg1 may bind to discrete regulatory regions of the *Atoh1* to temporally control its initial activation in Sox2^+^ progenitors and upregulation during HC fate commitment. Brg1-enrichment at the *Prox1* locus but not *p75* locus further indicates the essential function of Brg1-based chromatin remodeling in the activation of transcriptional programs in distinct subtypes of precursor cells during sensory epithelium development (Figure 7-figure supplement 7). Future work is necessary to elucidate how Brg1 interacts with other factors, including Eya1, Six1 and Sox2, to target its binding sites at the *Sox2*, *p27*^*Kip1*^ and *Atoh1* loci identified from our ChIP-seq data using a combination of transgenic, genomic and molecular approaches.

Our results suggest that Brg1-BAFs may have cell type-specific and context-dependent roles in inner ear development. During development, distinct BAF subunits interact with specific TFs, recruiting the complex to the sites of action of the corresponding factors to control distinct transcriptional programs (Ho et al., 2009; Lessard et al., 2007; Wang et al., 1996). Simultaneous interactions of Eya1/Six1 with BAF60a/c and BAF170/155 suggests formation of separate SWI/SNF complexes containing distinct BAF subunits in different subsets of Eya1- or Six1- expressing cells, in response to differentiation cues, that activate a diverse program of lineage-specific gene expression. This is in an agreement with the differential and overlapping patterns of BAF170/155/60a/c in the cochlear sensory epithelium. As BAF60a or BAF60c acts in recruiting TFs (Chen et al., 2012; Oh et al., 2008) and nuclear receptors (Hsiao et al., 2003) to the SWI/SNF chromatin remodeling complex, the variants a/c of the BAF60 structural subunit may recruit Eya1-Six1 to the SWI/SNF complexes to regulate stage-dependent and cell type-specific programs. In response to signals, the alternative usage of specific SWI/SNF variants by the prosensory progenitors may direct distinct differentiation programs. The selective assembly of SWI/SNF with BAF60c and BAF170 may activate HC-specific genes, while Brg1 along with BAF60a-Eya1 and BAF60a-Six1 may promote both HC and SC gene activation programs during terminal differentiation. The observation of a unique pattern of BAF170 expression in the pillar cells suggests that distinct subtypes of SCs may also require cell type-specific SWI/SNF complexes. We speculate that during otic development and differentiation, otic-specific activators may both recruit and require chromatin-remodeling activities for stable binding to the regulatory regions of otic- and neurosensory-specific genes. Future molecular studies will be required to elucidate if these complexes are recruited by transient interactions with different TFs including Eya1, Six1, and Sox2, and whether association of Eya1-Six1 with BAF60a/c recruits Eya1-Six1 into a Brg1-based SWI/SNF complex that is able to remodel chromatin and activate neurosensory-specific gene transcription.

In summary, our results indicate that Brg1 activity is required for *Eya1*/*Six1* expression and that Eya1-Six1 may use chromatin remodeling properties of the Brg1-BAFs to fine-tune control of neuronal or sensory cell fate determination. Activation of *Sox2* in the otic ectoderm requires both Eya1/Six1 function. However, in vivo, Sox2 alone without Brg1 activity or Eya1/Six1 activity is insufficient to induce *p27*^*Kip1*^ or *Atoh1* expression in the Sox2^+^ progenitors in the cochlea. These findings illustrate how chromatin-remodeling enzymes are intricately involved in the transcriptional control of the master regulators of development and the differentiation of neurosensory cells in the inner ear. Since ChIP-seq failed to recover Brg1 enrichment at *Neurog1* or *Neurod1* loci, how chromatin remodeling of the Brg1-BAFs intersects with chromatin remodeling of these neuronal cell-specific bHLH TFs remains to be demonstrated. In conclusion, the genome-wide maps of in vivo Brg1 binding sites generated in this study provide a foundation for understanding cochlea-specific genome-wide changes in chromatin structure driven by this key chromatin remodeling factor, which is critical to understanding the role of Brg1 in development and disease. This study provides not only a novel framework for understanding how TFs might direct neurosensory cell commitment and differentiation but also implies possibilities for therapeutic intervention by targeting the SWI/SNF complex in regeneration.

## Methods

### Animals, genotyping and tamoxifen administration

All animal protocols were approved by Animal Care and Use Committee of the Icahn School of Medicine at Mount Sinai (protocol #06-0807).

*Brg1*^*fl*^ (Sumi-Ichinose et al., 1997), *Eya1-CreERT2* (Xu et al., 2017; Xu et al., 2014), *Sox2-CreERT2* (Arnold et al., 2011), and *Eya1*^+/−^*;Six1*^+/−^ (Zheng et al., 2003) mice were maintained on a 129/Sv and C57BL/6J mixed background at the Icahn School of Medicine at Mount Sinai Animal Facility. Mice were bred using timed mating, and noon on the day of vaginal plug detection was considered as E0.5. For induction of the CreERT2 protein, tamoxifen (Sigma, T5648) was dissolved in corn oil (Sigma, C8267) and administrated (1.5 mg/10 g body weight) by oral gavage. Observed variations among *Brg1* mutants is likely due to pre-existing developmental variation between embryos when tamoxifen was given.

### Histology, immunostaining, and in situ hybridization (ISH)

Histological examination, whole-mount and section immunostaining and ISH were carried out according to standard procedures. Briefly, inner ears were fixed in 4% paraformaldehyde (PFA) for 1 hr at 4°C, dehydrated, and embedded in wax or OCT. Paraffin or frozen sections were generated at 6 μm. For ISH, tissues were fixed overnight. We used five or six embryos for each genotype at each stage for each probe and the result was consistent in each embryo.

For immunostaining, sections were stained with primary antibodies listed below. Cy2-, Cy5- and FITC-conjugated secondary antibodies were used. Hoechst 3342 was used for nuclear staining.

Primary antibodies: anti-Brg1 (ab110641, Abcam), -BAF170 (sc-17838, Santa Cruz), -BAF155 (sc-48350, Santa cruz), -BAF60a (sc-135843, Santa Cruz), -BAF60b (sc-101162, Santa Cruz), -BAF60c (ab171075, Abcam), -Sox2 (PA1-094, Thermo Fisher), -Myo7a (25-6790, Proteus and 138-1-s, DSHB), -p27^kip1^ (554069, BD Pharmingen), -Calretinin (MA5-14540, Thermo Fisher), -p75^NTR^ (#07-476, EMD Millipore), -S100A (ab11428, Abcam), -GLAST (ab416, Abcam), -Prox1 (AB5475, Millipore), -Brg1 (sc-10768, Santa Cruz), -BAF170 (sc-166237, Santa Cruz), Cy3-, Cy2-, Cy5- and FITC-conjugated secondary antibodies were used. Hoechst 3342 was used for nuclear staining.

### EdU and TUNEL Assays

The EdU assay was performed using a kit (catalog no. C10640, Life Technologies) following the manufacturer’s instructions. EdU was co-injected with tamoxifen at 9 am of E11.5 and embryos were harvested at afternoon of E13.5. EdU was also injected at 9 am of E9.5 embryos following tamoxifen treatment at 9 am of E8.0-8.5 and embryos were harvested 2 hours after EdU injection at noon of E9.5. The TUNEL assay was performed using the Apop Tag kit for in situ apoptosis fluorescein detection (catalog no. NC9815837, Millipore) following the manufacturer’s instructions.

### Cell counts and calibration

EdU-incorporated cells and Hoechst-stained nuclei in E9.5 otocyst were counted respectively. EdU-incorporated Sox2^+^ prosensory progenitors in the floor of the cochlear epithelium were counted in the entire cochlea. Values represent average number of EdU^+^ cells or proliferation rate of EdU^+^ cells/total Hoechst^+^ cells (±standard deviations) per otocyst or EdU^+^Sox2^+^ cells (±standard deviations) per section (6 μm) or per cochlea. 15 sections per cochlea and 6 cochleae for each sample were measured. Two-tailed Student’s *t*-test was used for statistical analysis.

### Co-immunoprecipitation

Embryonic cochleae (E13.5-14.5) were homogenized and lysed in 10 mM HEPES, pH7.5, 1.5 mM MgCl2, 10 mM KCl, 1 mM dithiothreitol (DTT) and protease and phosphatase inhibitors cocktail. After removal of cytoplasmic fraction, the crude nuclei pellet was lysed in nuclear extraction buffer (20 mM mM HEPES, pH7.5, 1.5 mM MgCl2, 420 mM NaCl, 0.2 mM EDTA, 25% glycerol, 1 mM dithiothreitol (DTT) and protease and phosphatase inhibitors cocktail). The extracted nuclear proteins were diluted in IP buffer (Tris-Cl, pH 8.0, 100 mM NaCl, 0.1% NP-40, 10% glycerol). The lysates were pre-cleared with protein A/G beads (sc-2003, Santa Cruz). After removal of the beads, the lysates were incubated with ~1 µg primary antibodies anti-Brg1 (ab110641, Abcam), anti-Flag (F1804, Sigma) or -anti-Six1 (HPA001893; Sigma) with rotation at 4 °C, overnight. Lysates were incubated with Protein A/G beads to precipitate complexes for 3 h at 4 °C with rotation. The immunocomplexes were recovered by centrifugation and washed three times with IP buffer. SDS buffer was added to the precipitate and boiled for 5 min. Proteins were size separated in SDS–PAGE. The gels were blotted onto an Immobilon-P membrane (IPVH00010, Millipore), blocked with 5% non-fat dry milk and incubated with the previously described antibodies. HRP-conjugated secondary antibodies (Goat anti-Mouse IgG (H+L), A4416, Sigma; Goat anti-Rabbit IgG (H+L), 31460, Thermo Scientific) were used for detection using the enhanced chemiluminescence (ECL) method (WBKLS0500, Millipore).

### ChIP-seq and ChIP-qPCR

Cochleae were dissected from E13.5-14.0 embryos and cross-linked with 1% formaldehyde at room temperature for 30 mins. Samples were then homogenized and lysed in cold lysis buffer (50 mM HEPES-KOH, pH 7.5, 140 mM NaCl, 1 mM EDTA, 10% glycerol, 0.5% NP-40, 0.25% Triton X-100, 1× protease inhibitors) and gently rocked at 4°C for 10 minutes in 15 ml conical tubes. Cells were pelleted at 2000 g at 4°C and resuspended in cold wash buffer (10 mM Tris-HCl, pH 8.0, 200 mM NaCl, 1 mM EDTA, 0.5 mM EGTA, 1× protease inhibitors) and gently rocked at 4°C for 10 minutes in 15 ml conical tubes. Cells were pelleted at 2000 g at 4°C in a table top centrifuge and resuspended in 1 ml cold sonication buffer (10 mM Tris-Cl, pH 8.0, 2 mM EDTA, 0.1% SDS) and sonicated to 200-500 bp fragments using a Covaris S220 Focused-ultrasonicator. Sonicated chromatin was cleared by pelleting insoluble material at 13,000 RPM at 4°C followed by preclear with protein A/G beads and incubation with 1~2 μg antibody overnight (anti-Brg1, ab110641, Abcam; anti H3K27ac, ab4792, Abcam). Next, Protein A/G beads (20 μl) were added to the lysate and incubated at 4°C for 5 hrs. Immunoprecipitated material was washed with low salt buffer (20 mM Tris-Cl 8.0, 150 mM NaCl, 2 mM EDTA, 1% Triton X-100, 0.1% SDS), high salt buffer (20 mM Tris-Cl pH 8.0, 500 mM NaCl, 2 mM EDTA, 1% Triton X-100, 0.1% SDS), and LiCl wash buffer (10 mM Tris-HCl pH 8.0, 250 mM LiCl, 1 mM EDTA, 1% NP-40, 1% sodium deoxycholate) and one time with TE plus NaCl, followed by elution and reverse crosslinking overnight at 65°C. The quality controls of ChIPed DNA was performed with Qubit 2.0 Fluoremeter using dsDNA HS assay Kit (Q32854, ThermoFisher Scientific) and Agilent 2200 TapeStation System using Hightivity D1000 Reagenets (5067-5585, Agilent). The libraries for sequencing were prepared with the ThruPlex DNA-seq Kit (R400429, Rubicon Genomics) and sequenced on Illumina HiSeq 2500 system.

For ChIP-qPCR, chromatin derived from embryonic otocysts or from E13.5 cochleae and IgG, anti-Brg1 (ab110641, Abcam), anti-BAF60a antibody (ab224229, Abcam), or anti-BAF60c (ab171075, Abcam) was used for ChIP respectively. The ChIPed DNAs were subjected to quantitative PCR (qPCR) amplification with StepOnePlus PCR system and SYBR green PCR Master Mix kit (4309155, Applied Biosystems). The enrichment fold of IP over mock IP was calculated using the comparative Ct (threshold cycle) method. Data was normalized with inputs and the enrichment of mock IP was considered 1-fold. This experiment was repeated three times and each qPCR was performed in triplicate. The primers used for ChIP-qPCR are listed in Supplementary Table 4. The DNA positions are denoted relative to the transcriptional start site (+1).

### Peak calling and gene otology analysis

The ChIP-seq data were first checked for quality using the various metrics generated by FastQC (v0.11.2) (http://www.bioinformatics.babraham.ac.uk/projects/fastqc). Raw sequencing reads were then aligned to the mouse mm10 genome using default settings of Bowtie (v2.2.0) (Langmead et al., 2009). Only uniquely-mapped reads were retained and duplicates were removed. Peak-calling was performed using MACS (v2.1.1) (Zhang et al., 2008) with various p-value cutoffs as reported in the main text. Motif enrichment analysis was performed using the Homer package (v4.8.3) (Heinz et al., 2010). The peak annotation and gene ontology analysis was performed using GREAT program (McLean et al., 2010) and Panther classification system (Thomas et al., 2003). The conservation between mouse and human genome sequences was analyzed online with ECR Brower (Ovcharenko et al., 2004).

### Accession numbers

The ChIP-seq data reported in this paper were submitted to GEO (GSE119545).

## Acknowledgements

We thank L. Zhang for technical assistance. This work was supported by the NIH RO1 DC014718 (PXX) and NYSTEM C029566 (PXX).

## Author contributions

J.X., J.L., Y.H.E.L, A.R., T.Z., H.J., data collection and analysis, validation, writing-review/editing; L.S., B.F., data analysis, writing-review/editing; P.X. X., conceptualization, supervision, funding acquisition, data analysis, writing-original draft, writing-review/editing. All authors approved the final version of the manuscript.

## Supplemental Figure legends

**Figure supplement 1.**
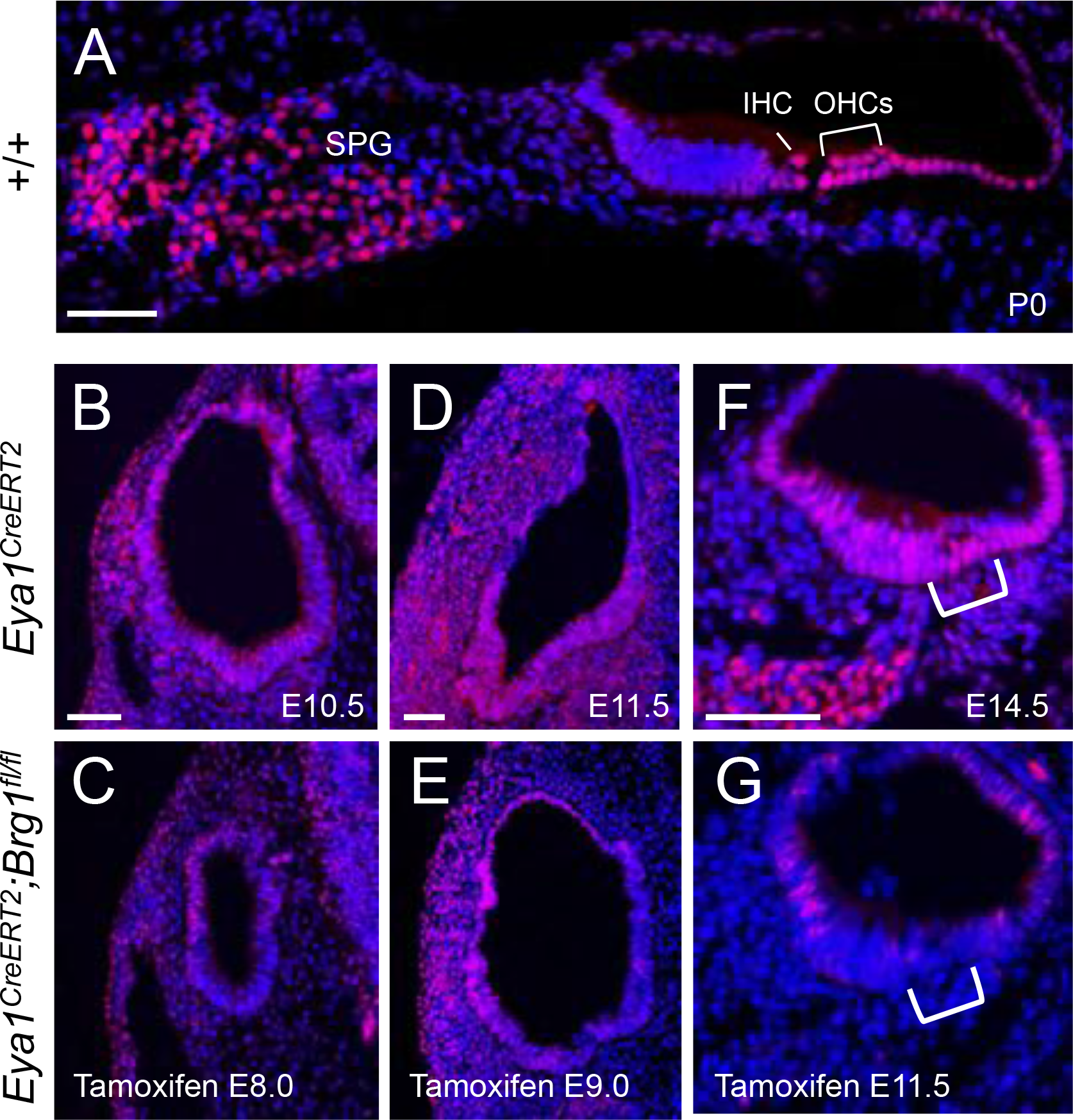
Immunostaining confirms selective deletion of Brg1 in the otocyst and cochlear epithelium using *Eya1*^*CreERT2*^. (A) Anti-Brg1 immunostaining on P0 cochlear section. SPG, spiral ganglion; IHC, inner hair cell; OHCs, outer hair cells. (B-G) Anti-Brg1 immunostaining on otic sections from *Eya1*^*CreERT2*^ (B,D,F) and *Eya1*^*CreERT2*^*;Brg1*^*fl/fl*^ littermate (C,E,G) embryos at E10.5 (given tamoxifen at E8.5-9.5; B,E), E11.5 (given tamoxifen at E9.0-E10.0; C,F) and E14.5 (given tamoxifen at E11.5-12.5; D,G) showing reduction of Brg1 in the ventral region of the mutant otocyst or in the primordial organ of Corti (bracket). As Eya1 is also strongly expressed in surrounding periotic mesenchyme, this also led to removal of Brg1 in Eya1-expressing periotic mesenchyme. (C,E, compare with B,D). Scale bars: 50 μm.

**Figure supplement 2.**
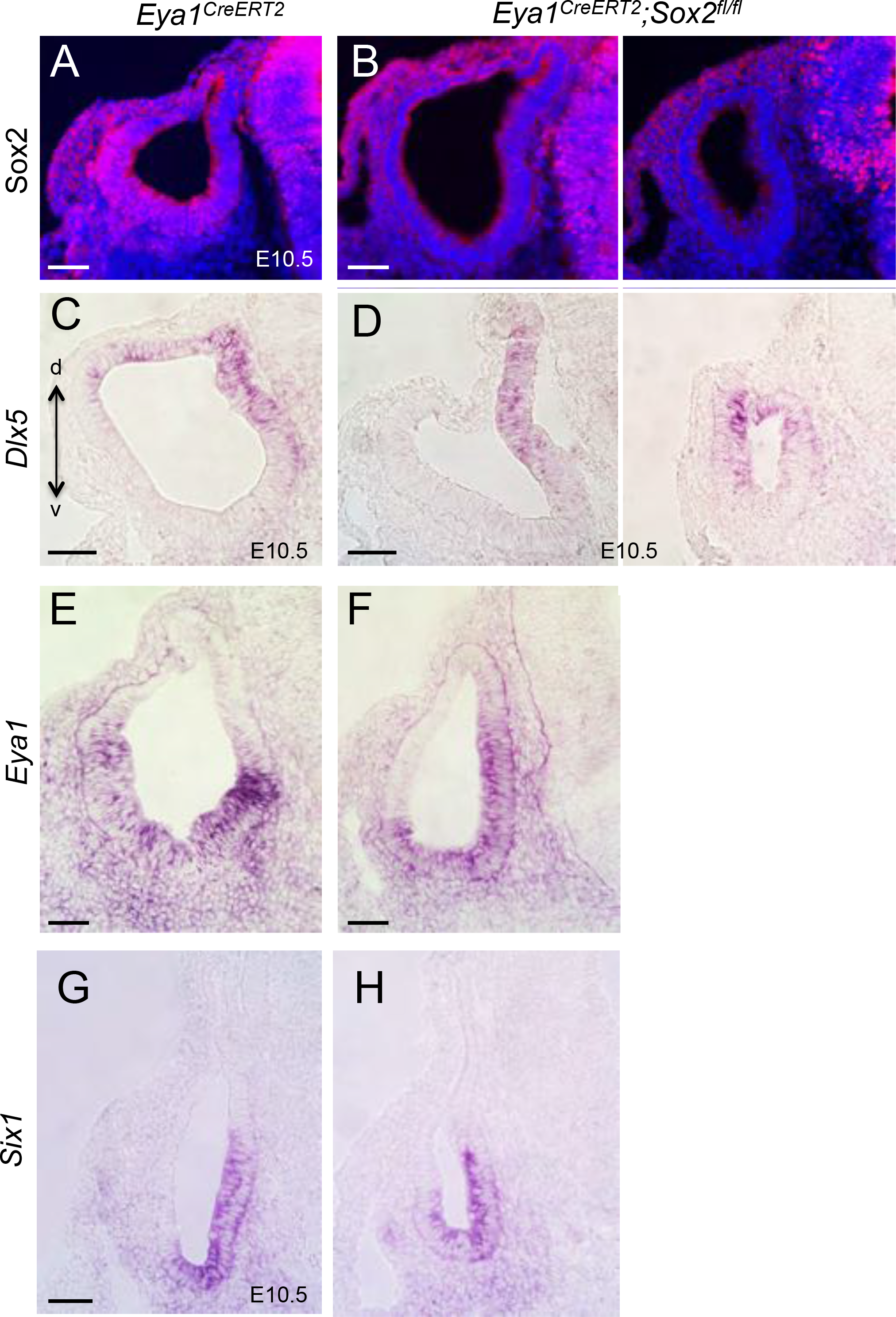
Conditional deletion of *Sox2* in the otic placode does not alter the pattern of *Dlx5*, *Eya1* and *Six1* expression. (A,B) Immunostaining for Sox2 confirms deletion of Sox2 in E10.5 otocyst using *Eya1*^*CreERT2*^ (tamoxifen given from E7.5-8.5). (C,D) In situ hybridization on transverse sections of *Eya1*^*CreERT2*^ and *Sox2*^*Cko/Cko*^ otocysts for *Dlx5* (C,D), *Eya1* (E,F) and *Six1* (G,H). Abb.: a, anterior; d, dorsal; p, posterior; v, ventral. Scale bars: 50 μm.

**Figure supplement 3.**
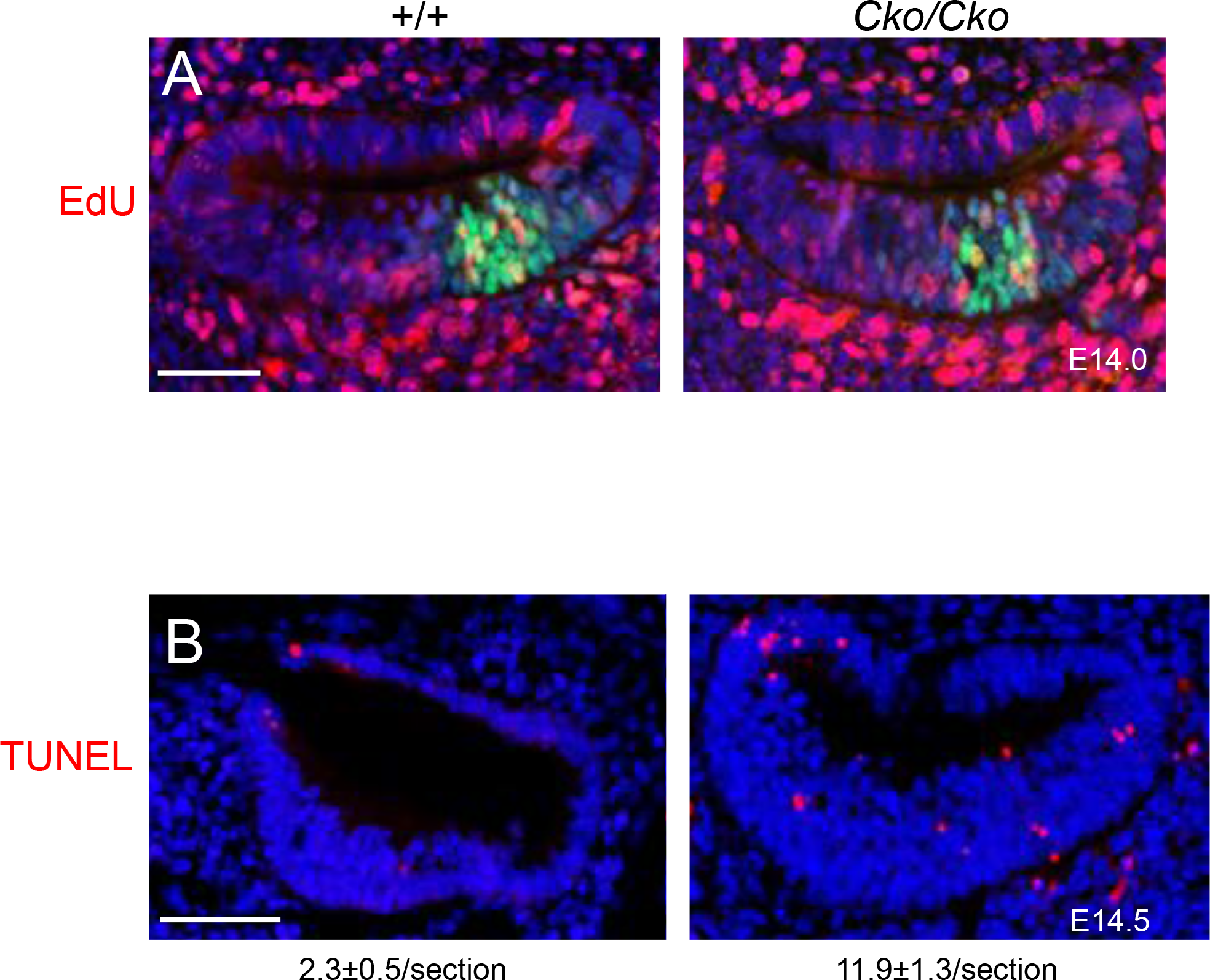
Brg1 is necessary for normal cell proliferation and survival in the cochlear sensory epithelium. (A) EdU-incorporation (red) in E13.5-14.0 wild-type and *Brg1*^*Cko/Cko*^ cochlea given tamoxifen and EdU at E11.5. Sections were co-immunostained with Sox2 (green). (B) TUNEL analysis of cochlea of wild-type and *Brg1*^*Cko/Cko*^ embryos. Statistical analysis of EdU-labeled Sox2^+^ cells or apoptotic cells from 3 embryos per genotype showing the average number (±standard deviations) of Sox2^+^/EdU^+^ cells or apoptotic cells per section (6 µm) or per cochlea. *P*=0.0028 for B. *P*-value was established for +/+ and *Brg1*^*Cko/Cko*^ using Two-tailed Student’s *t*-test. Scale bars: 50 μm.

**Figure supplement 4.**
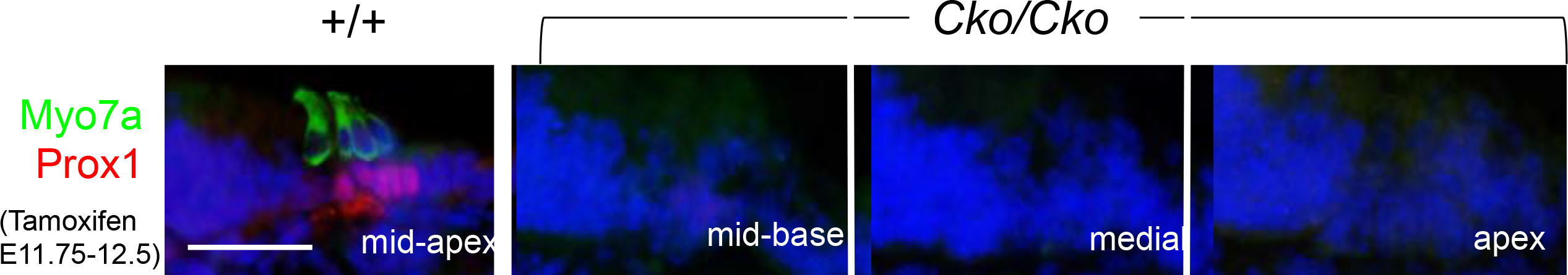
Disruption of hair and supporting cell differentiation in *Brg1*^*Cko/Cko*^ cochlea. Immunostaining for Myo7a (green) and Prox1 (red) on cochlear sections from E18.5 inner ears of wild-type and *Brg1*^*Cko/Cko*^ littermates (given tamoxifen at E11.5-12.5. Scale bars: 30 μm.

**Figure supplement 5.**
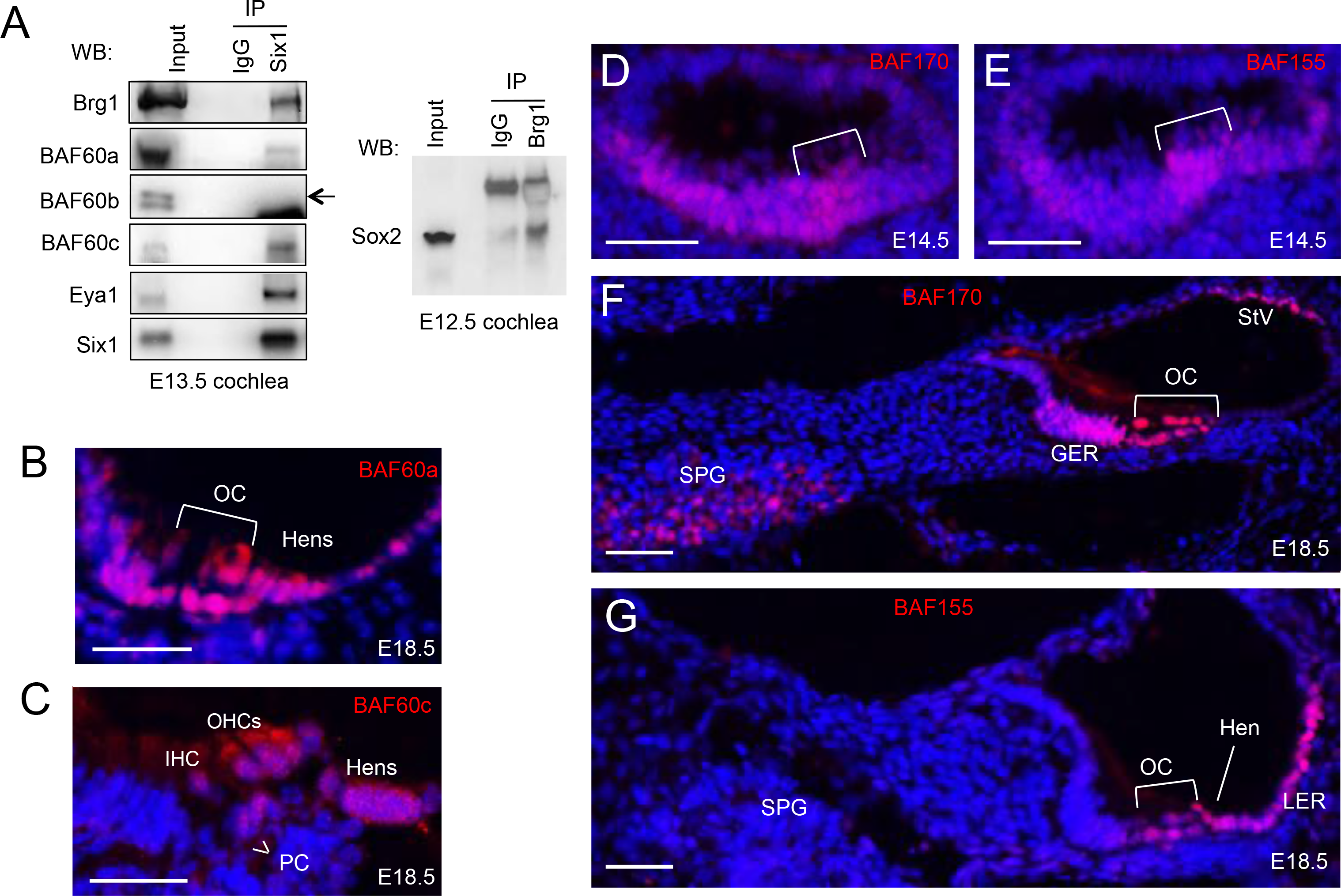
Brg1 physically interacts with Six1 and Sox2 and differential and overlapping patterns of BAF60a/c, BAF170 and BAF155 expression in the developing cochlea. (A) CoIP analysis. Nuclear extracts were prepared from E13.5 or E12.5 cochleae. Antibodies for IP and western blot are indicated. Input was 5% of the amount used for IP. Arrow indicates no interaction between BAF60b and Six1. (B,C) Immunostaining for BAF60a (B) or BAF60c (C) on cochlear sections of wild-type embryo at E18.5. (D,E) Immunostaining for BAF170 (D,F) or BAF155 (E,G) on cochlear sections of wild-type embryo at E14.5 or E18.5. Brackets mark the primordial organ of Corti. SPG, spiral ganglion. Scale bars: 30 μm (B,C), 50 μm (D-G).

**Figure supplement 6.**
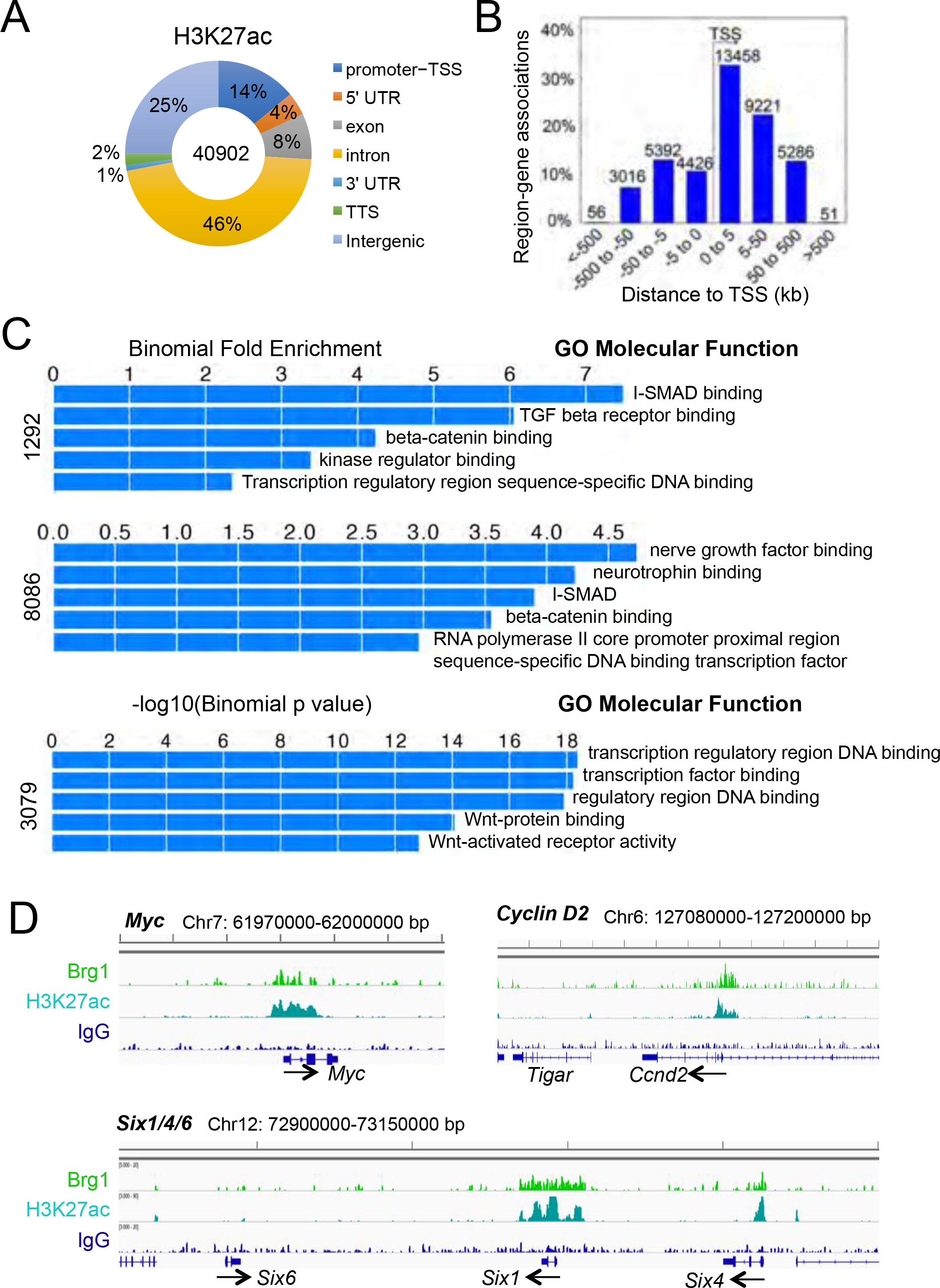
Genomic binding features of Brg1 and H3K27ac in E13.5 cochlea. (A) Genomic distribution of H3K27ac-enriched regions. Peaks were called with the MACS program. (B) Distribution of H3K27ac peaks relative to TSSs. (C) Differential enrichment for functional annotation terms associated Brg1-enriched regions. Shown are the top 5 enriched “molecular functions” associated with the 1292 or 8086 Brg1-enriched regions or specific to the 3079 Brg1 regions with H3K27ac association. (D) Genomic view showing Brg1 enrichment at *Myc, Cyclin D2 and Six1/4* but not at *Six6* locus.

**Figure supplement 7.**
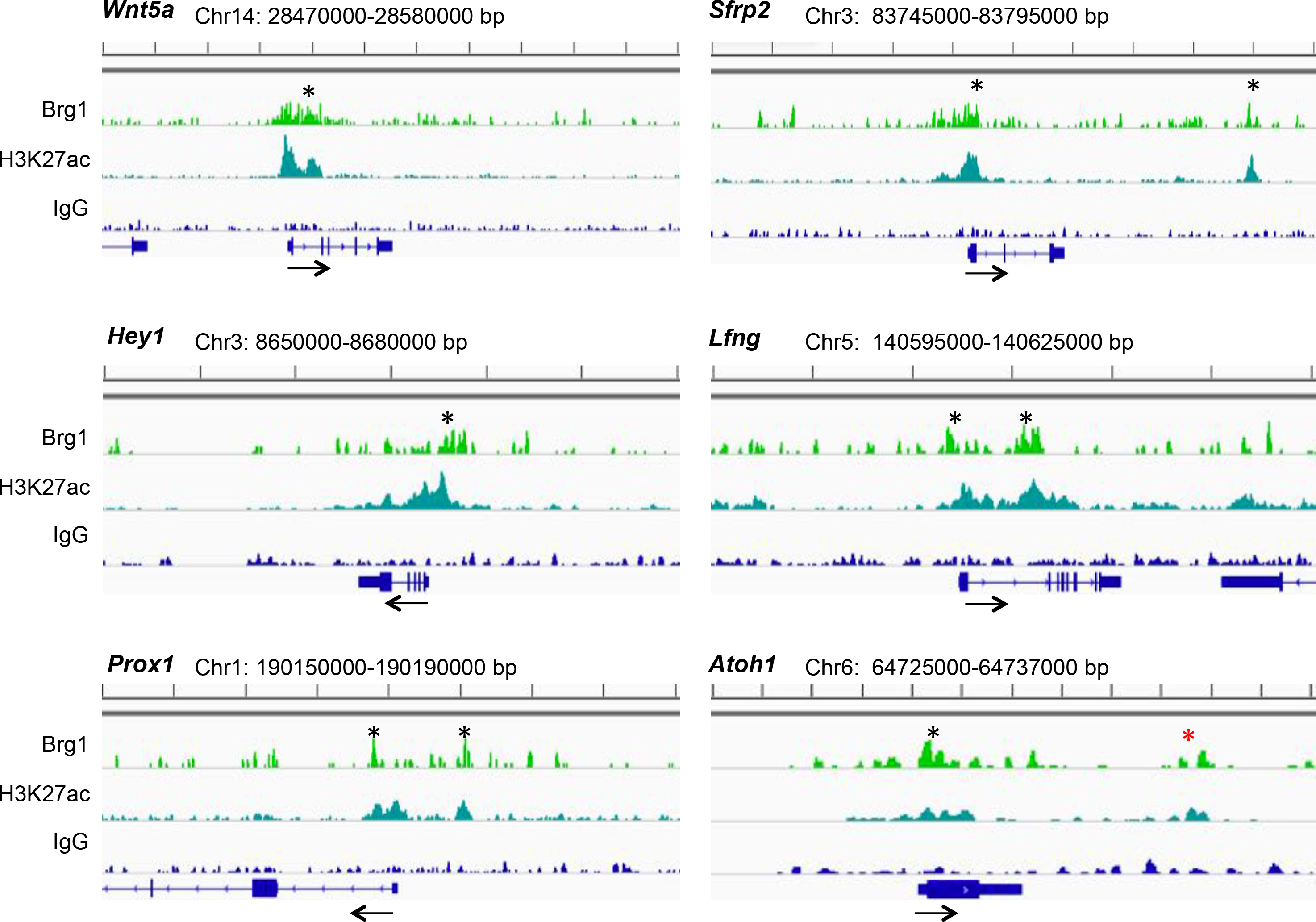
Brg1 associates to both proximal and distal sites of key loci. Genomic view showing Brg1 enrichment at genes involved in different pathways regulating sensory cell development. *Wnt5a* and *Sfrp2*—involved in Wnt-signaling, *Hey1* and *Lfng*—involved in Notch signaling, *Prox1*—a marker specific for supporting cells; *Atoh1*—critical for hair cell differentiation.

**Figure supplement 8.**
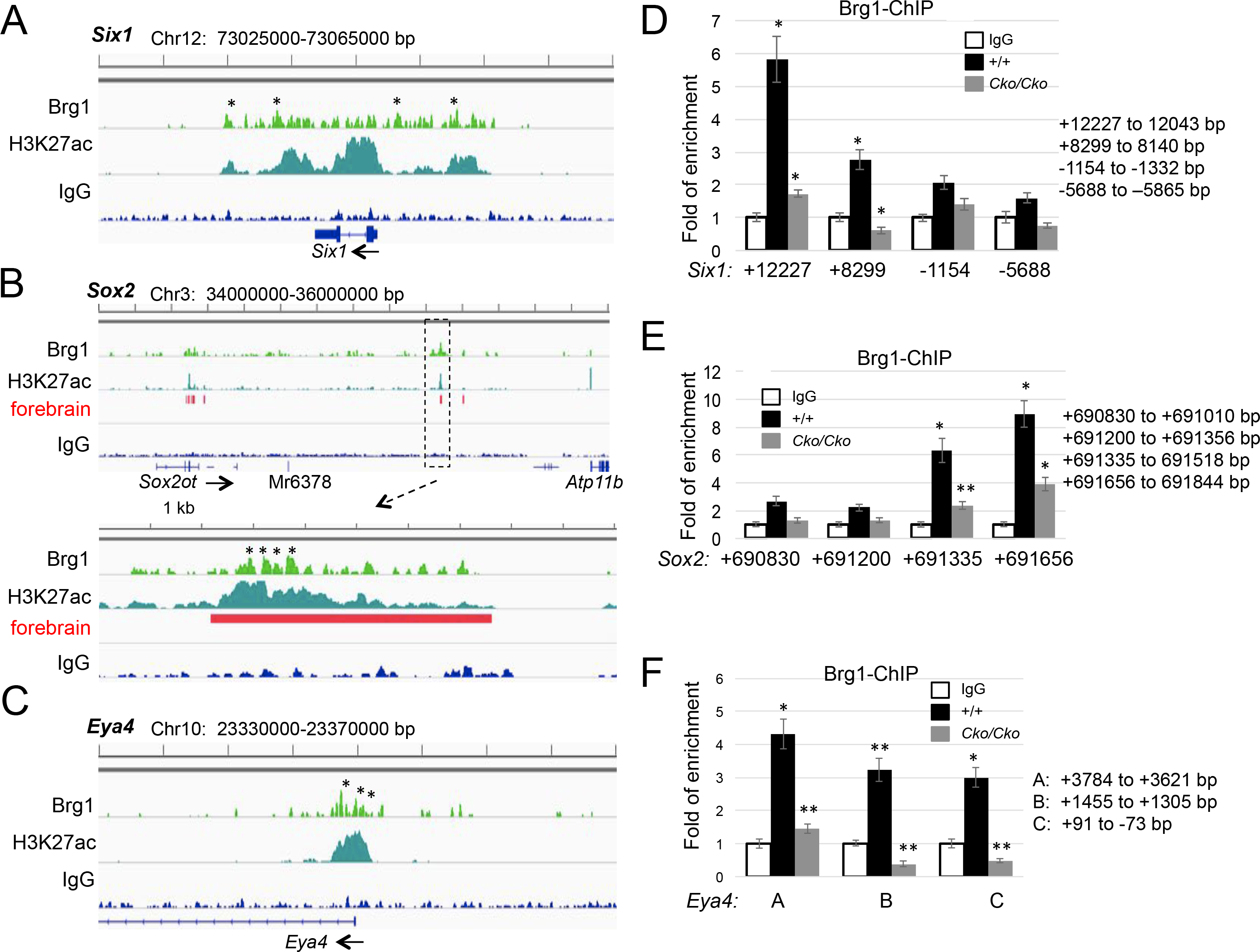
Brg1-bound regions at the *Six1*, *Sox2* and *Eya4* loci and ChIP assays confirm Brg1 enrichment at these loci. (A-C) Genomic view showing Brg1-bound regions at distal regions of the *Six1, Sox2* and *Eya4* loci. Regions selected for ChIP assays (corresponding to the Brg1 signals marked by asterisks) for each gene are indicated. At the *Sox2* locus, four sites within an intergenic Brg1-bound region (located ~691 kb downstream of *Sox2*; boxed by dashed line) were selected for ChIP assays. This region was also recovered in the Brg1 ChIP-seq on E11.5 forbrain (Attanasio et al., 2014) (red bar). Primers for amplifying each element are listed on Supplementary file 4. (D-F) ChIP-qPCR data using chromatin derived from otocysts of wild-type or *Brg1*^*Cko/Cko*^ littermates at E10.5 (given tamoxifen at E8.0-9.0) and anti-Brg1. Representative of three independent experiments, n=3 for each group, \**P*<0.01, \*\**P*<0.05 determined by Student’s t-test).

